# Development of photosynthetic carbon fixation model using multi-excitation wavelength fast repetition rate fluorometry in Lake Biwa

**DOI:** 10.1101/2020.08.10.244012

**Authors:** Takehiro Kazama, Kazuhide Hayakawa, Victor S. Kuwahara, Koichi Shimotori, Akio Imai, Kazuhiro Komatsu

## Abstract

Direct measurements of gross primary productivity (GPP) in the water column are essential, but can be spatially and temporally restrictive. Fast repetition rate fluorometry (FRRf) is a bio-optical technique based on chlorophyll *a* (Chl-*a*) fluorescence that can estimate the electron transport rate (ETR_PSII_) at photosystem II (PSII) of phytoplankton in real time. However, derivation of phytoplankton GPP in carbon units from ETR_PSII_ remains challenging because the electron requirement for carbon fixation (Ф_e,C_) can vary depending on multiple factors. Also, the FRRf is still relatively novel, especially in freshwater ecosystems where phosphorus limitation and cyanobacterial blooms are common. The goal of the present study is to construct a robust Ф_e,C_ model for freshwater ecosystems using simultaneous measurements of ETR_PSII_ by FRRf with multi-excitation wavelengths coupled with traditional carbon fixation rate by the ^13^C method. The study was conducted in oligotrophic and mesotrophic areas in Lake Biwa from July 2018 to May 2019. The combination of excitation light at 444, 512 and 633 nm correctly estimated ETR_PSII_ of cyanobacteria. The range of Ф_e,C_ in the phytoplankton community varied from 1.1 to 31.0 mol e^−^ mol C^−1^ during the study period. Generalized liner model showed the best model including 12 physicochemical and biological factors explained 67% of the variance in Ф_e,C_. Among all factors, water temperature was the most significant, while PAR intensity was not. The GPP values estimated by FRRf (*GPP_f_*) with the best Ф_e,C_ model relative to ^13^C (*GPP_13C_*) varied 0.5–1.5. Further, *GPP_f_* estimated with more parsimonious Ф_e,C_ models were also comparable to *GPP_13C_*. This study quantifies the applicability of the *in situ* FRRf methodology, and supports continuous monitoring of GPP by FRRf in lakes with large spatio-temporal variability of environmental conditions and phytoplankton assemblages.

## Introduction

Phytoplankton are the most important primary producers in the aquatic food web [1]. Changes in phytoplankton primary productivity can affect the food chain length [2,3], material cycles [4,5], and biomass of higher trophic organisms [6–8]. Phytoplankton community productivity is affected and must rapidly adapt to various environmental factors [9–11] due to rapid growth rates and short generation times [12]. In order to evaluate the effect of variability in environmental factors on aquatic communities and ecosystems, continuous observation of phytoplankton primary productivity is necessary [4,7,13].

Traditional chemical methods of primary production measurements, *i.e.*, the ^14^C method [14], the ^13^C method [15,16], the light-dark bottle method [17], and the ^18^O method [18], require handling radioisotope (^14^C) and/or incubation time for several hours. Thus, primary production studies using these techniques can be limiting when trying to assess temporal and spatial variability. Fast repetition rate fluorometry (FRRf; Table 1), a chlorophyll *a* fluorescence-based method, has been developed as an advanced bio-optical technique for real-time measurement of phytoplankton primary productivity, mainly in marine ecosystems [19–27]. The FRRf method enables induction and measurement of a range of chlorophyll *a* fluorescence yields and parameters specific to photosystem II (PSII) [19,20,28], and, in turn, enables estimation of the *in vivo* electron transport rate in PSII (ETR_PSII_) and gross primary productivity (GPP) by theoretical models of photosynthesis [19,28,29].

**Table 1.**
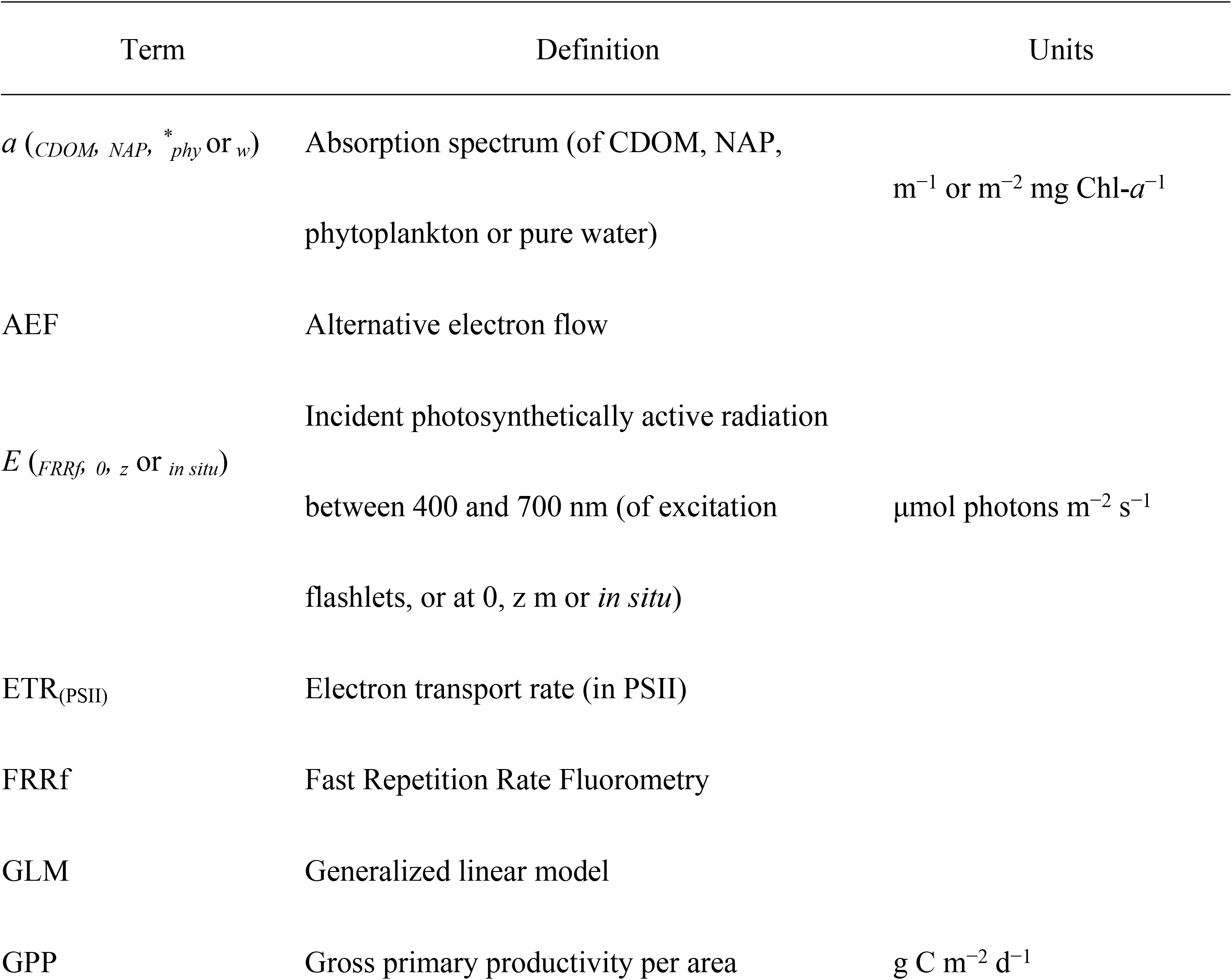
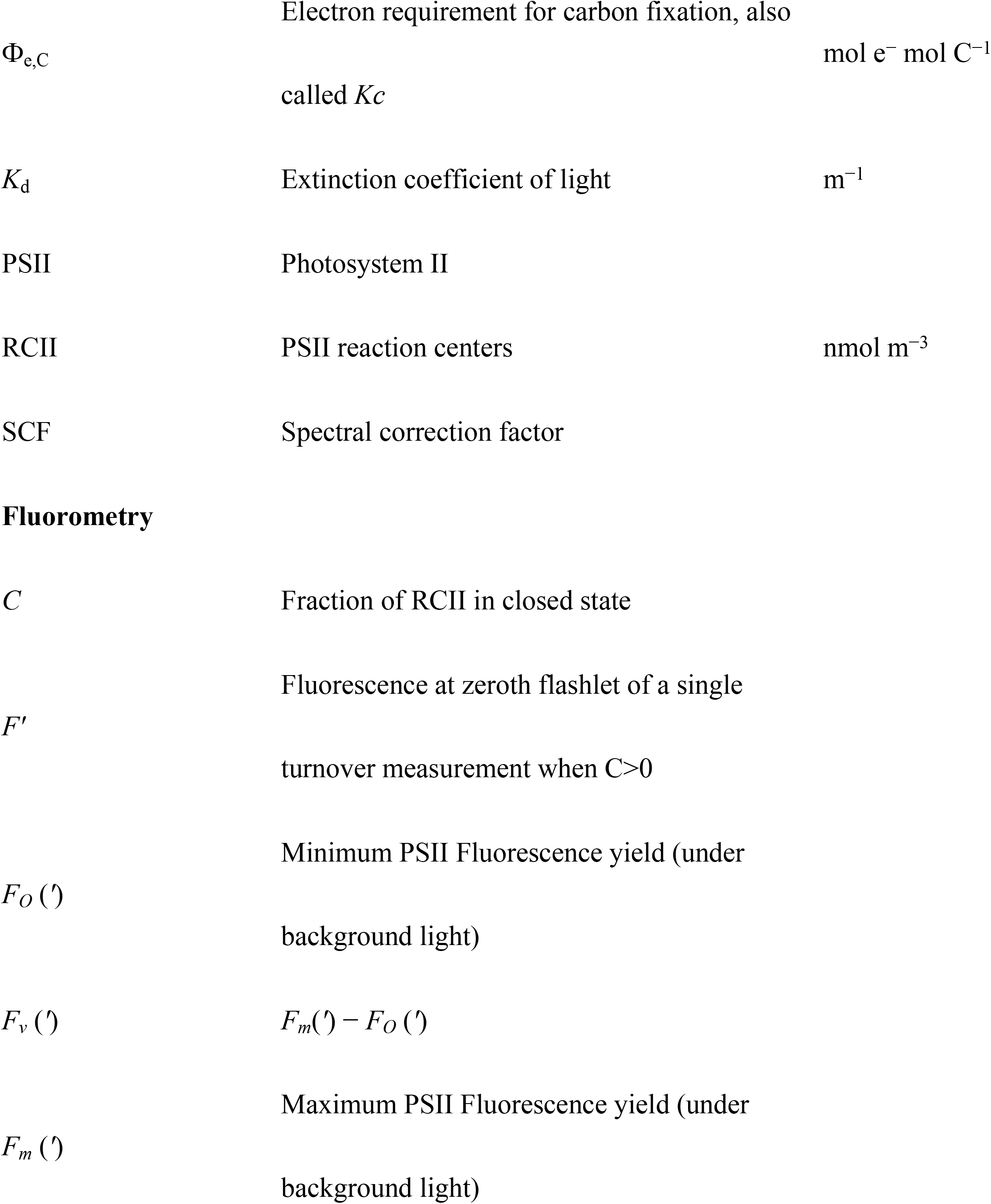
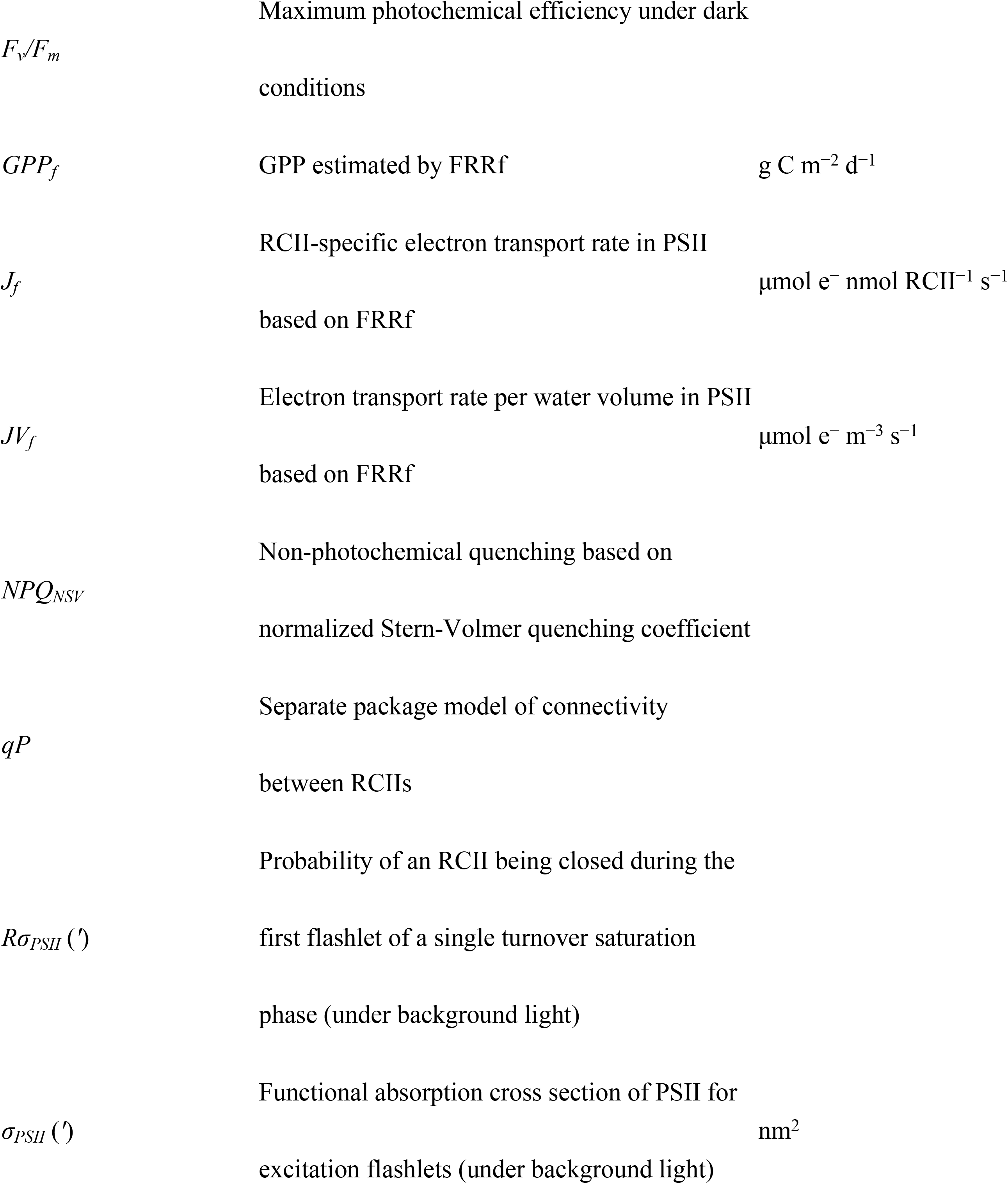
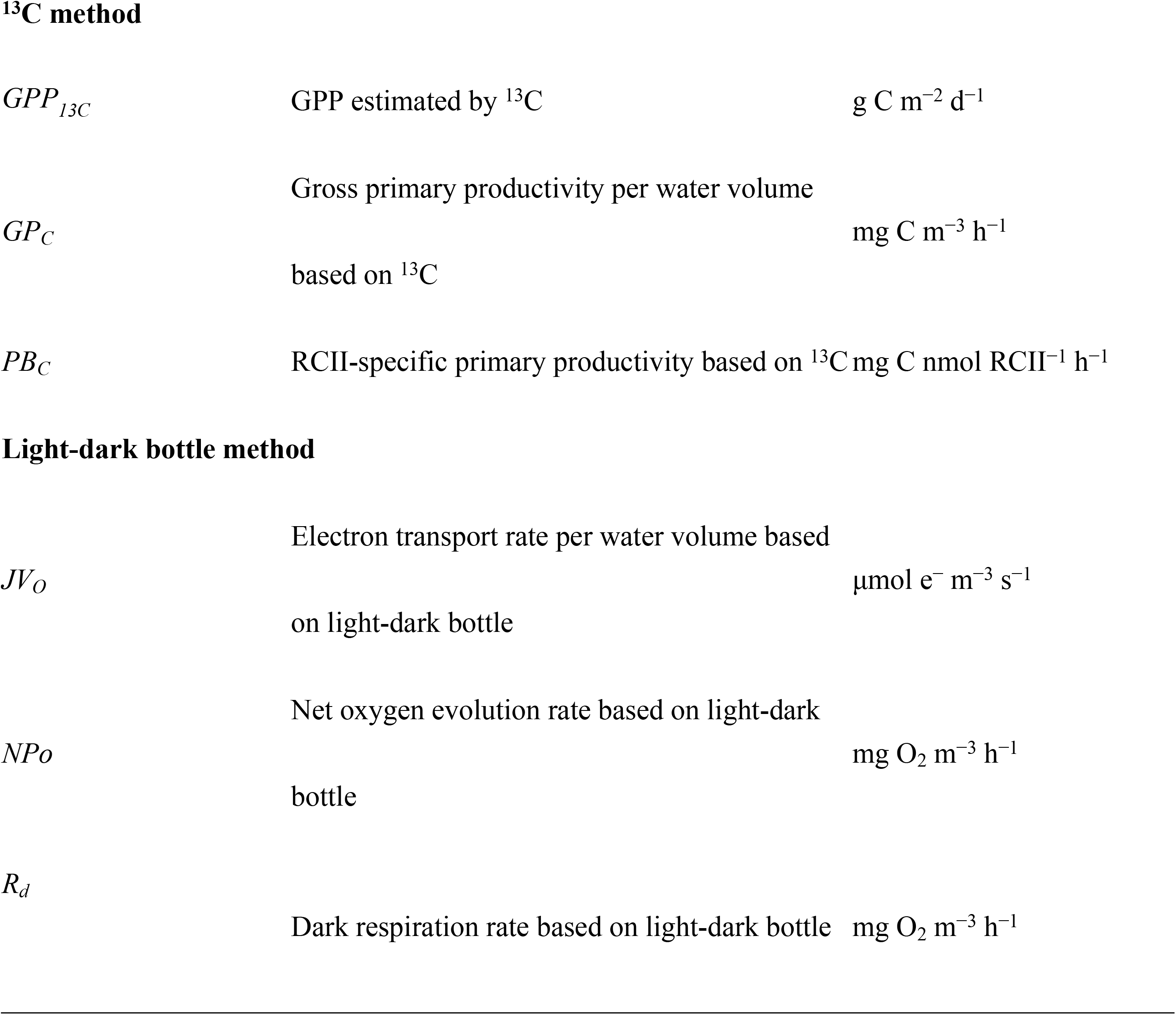
Terms used within this manuscript.

Previous studies demonstrated that GPP estimated from FRRf measurements correlated well with results from conventional methods, including the ^13^C method [25,26,30,31] and the light-dark bottle method [24]. However, FRRf tended to over- and under-estimate GPP compared to the ^14^C and ^13^C methods, and the light-dark bottle and ^18^O methods, respectively [32]. These discrepancies in GPP measurements are dependent on the targeted products, *i.e.* oxygen or particulate organic carbon, in the photosynthesis cycle [33]. In order to account for the measurement discrepancies, recent studies are examining the electron requirement for carbon fixation (Ф_e,C_, also called *Kc*) by comparing the FRRf-derived ETR_PSII_ to the GPP rate by traditional methods [25–27,34]. The Ф_e,C_ is affected by multiple spatio-temporal variations in physical and chemical factors [35–40], and by phytoplankton community composition [26,41–44]. For example, Ф_e,C_ is relatively higher in the open ocean compared to coastal areas due to differences in the light environment conditions; light availability is higher in open oceans [35]. More specifically, excess light energy enhances photo-oxidative damage and alternative electron transport such as the Mehler reaction that can increase Ф_e,C_ [45]. In addition to ambient light conditions, nutrients can also play an important role in determining Ф_e,C_ of the phytoplankton community [27,39]. For example, Schuback et al. [27] described the negative relationships between Ф_e,C_ and nitrate concentrations in the Arctic Ocean suggesting the variable effects of nutrient availability. The multitude of interacting factors affecting the value of Ф_e,C_ for converting the ETR_PSII_ to GPP make it difficult to establish a general model applicable to different ecosystems [35]. Therefore, in order to construct a robust ETR_PSII_-GPP model, it is necessary to accumulate FRRf and its corresponding GPP data in various aquatic environments.

In terms of physicochemical and biological conditions, freshwater ecosystems differ considerably from marine ecosystems. For example, cyanobacteria (blue-green algae) can frequently form dense surface blooms in meso–eutrophic lakes [46,47]. Previous studies have suggested cyanobacterial blooms can significantly affect ETR_PSII_ measurements due to spectral mismatch between FRRf excitation wavelengths and the absorption spectrum of cyanobacteria [21,26,36,41,48]. For example, Raateoja et al. [41] found that filamentous cyanobacteria *Nodularia spumigena* and *Aphanizomenon* sp. had absorption peaks around 630 nm, and the photosynthetic activity of these species could not be measured by FRRf with an excitation light around 475 nm (targeting Chl-*a*). Cyanobacteria have multiple absorption peaks around 500–570 nm and 630 nm based on antenna pigments [49]. Thus, it is critical that the FRRf excitation wavelengths correspond to the absorption spectrum of cyanobacteria (or the dominant group) for accurate estimation of primary productivity and model development in freshwater ecosystems [21,41,50].

Phytoplankton primary productivity is also more likely to be phosphorus-limited in freshwater ecosystems [51,52], while more likely to be nitrogen-limited in marine environments [53,54]. Whereas nitrogen limitation depresses cellular Chl-*a* concentration, phosphorus limitation inhibits ATP synthesis, which can affect photochemical energy conversion in algae [55]. In fact, Ф_e,C_ of marine phytoplankton increases under nitrogen limitation [27,39]. Regrettably, phosphorus stoichiometry influence of Ф_e,C_ in freshwater phytoplankton remains unknown, and FRRf studies in freshwater environments in general are still limited [56–58]. Due to the differences between marine and freshwater ecosystems, it is essential to employ suitable excitation wavelength combinations and measure phosphorus concentration for correctly estimating both the ETR_PSII_ and GPP of freshwater phytoplankton communities.

The goal of the present study is to construct a robust Ф_e,C_ model applicable to freshwater ecosystems using simultaneous measurements of ETR_PSII_ by FRRf coupled with traditional carbon fixation rate by the ^13^C method in Lake Biwa. From this context, we first evaluated the performance of FRRf with multi-excitation wavelengths (444, 512, and 633 nm; S1 Appendix A) during cyanobacterial blooms. Then, we evaluated the relative importance of variable environmental and biological factors, including phosphorus concentration, by determining Ф_e,C_ by statistical models with multiple variables. This study shows the utility of *in situ* FRRf measurements using excitation wavelength 633 nm during cyanobacterial blooms, and the extent to which physicochemical factors and phytoplankton community composition influence Ф_e,C_ estimation.

## Materials and methods

### Study site

The study was conducted at Lake Biwa (670 m^2^ surface area with a mean depth of 43 m) on Honshu Island, Japan (Fig. 1). The North Basin is a deep, oligotrophic area, while the South Basin is a shallow, mesotrophic area [59]. Phytoplankton communities are substantially different between the two basins, especially in terms of cyanobacteria abundance [60]. Sampling was carried out at the long-term survey stations, 12B (62 m depth) and 17B (89 m depth) in the North Basin, and at 6B (4 m depth) and 9B (5.6 m depth) in the South Basin, from July 2018 to May 2019 (Table 2).

**Fig. 1.**
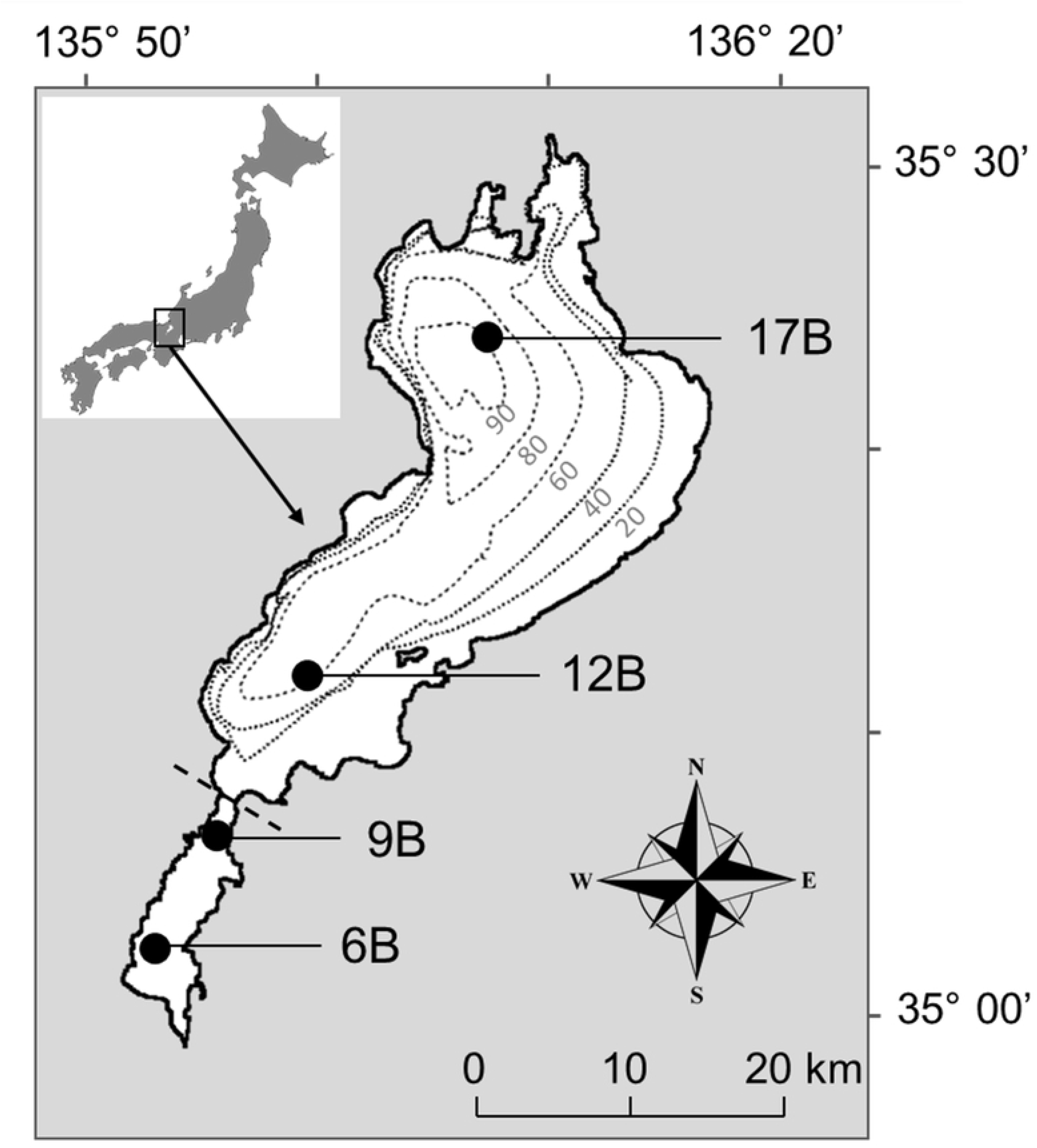
Map of study sites in Lake Biwa, Japan. Stations 6B and 9B represent the South Basin, while Stations 12B and 17B were selected as representatives of the North Basin. Grey dotted lines indicate the isobaths (in m), and dashed line represents the boundary of the basin.

**Table 2.**
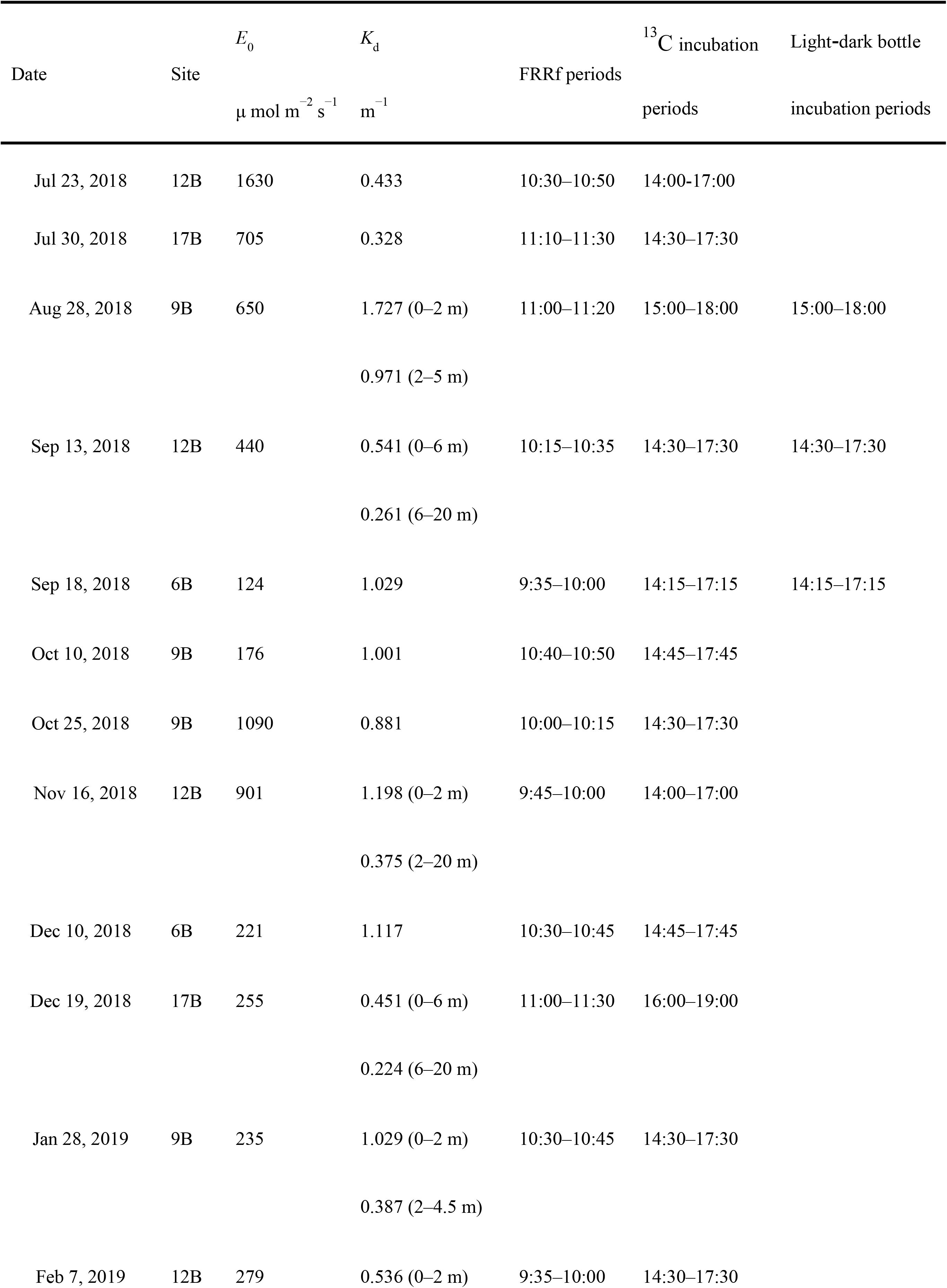
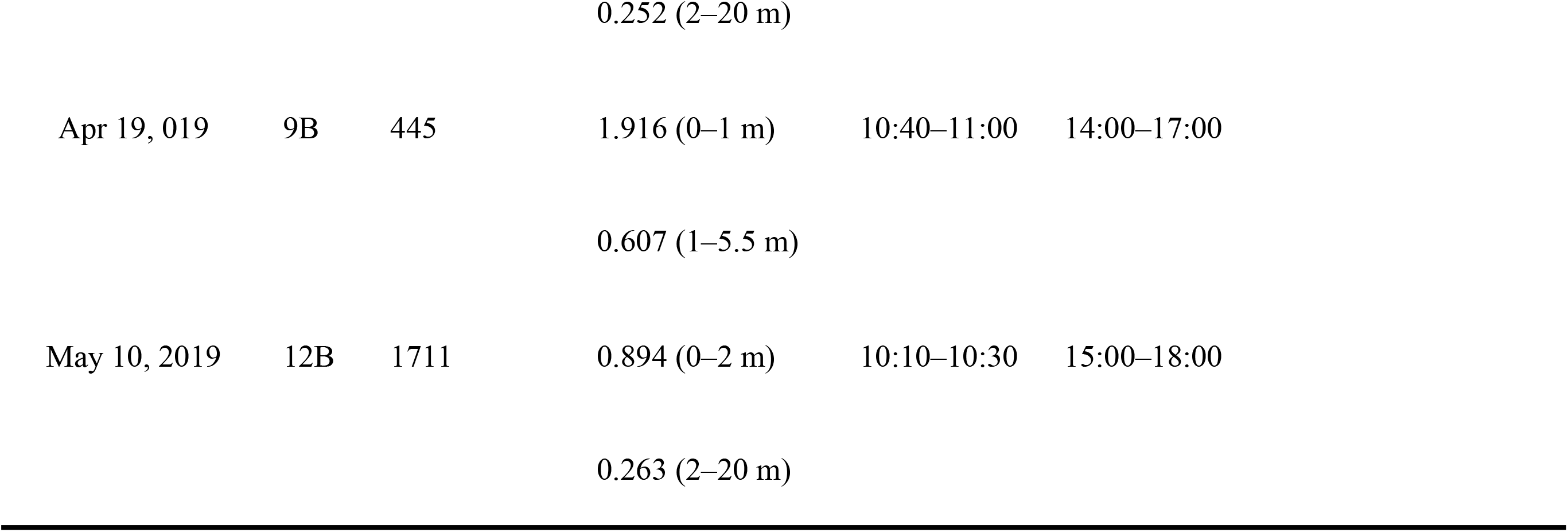
Sampling date, Station ID, light environment (*E*_0_ and *K*_d_), and time periods for FRRf and ^13^C incubation on each sampling date. *E*_0_: sub-surface PAR at 0 m at the time of FRRf sampling; *K*_d_: diffuse attenuation coefficient of PAR calculated from equation (1). Additional calculations of *K*_d_ were made for each layer when the logarithmic slope significantly changed with depth.

| Date | Site | $E_0$<br>$\mu\text{ mol m}^{-2}\text{ s}^{-1}$ | $K_d$<br>$\text{m}^{-1}$ | FRRf periods | <sup>13</sup>C incubation<br>periods | Light-dark bottle<br>incubation periods |
| --- | --- | --- | --- | --- | --- | --- |
| Jul 23, 2018 | 12B | 1630 | 0.433 | 10:30–10:50 | 14:00–17:00 |  |
| Jul 30, 2018 | 17B | 705 | 0.328 | 11:10–11:30 | 14:30–17:30 |  |
| Aug 28, 2018 | 9B | 650 | 1.727 (0–2 m) | 11:00–11:20 | 15:00–18:00 | 15:00–18:00 |
|  |  |  | 0.971 (2–5 m) |  |  |  |
| Sep 13, 2018 | 12B | 440 | 0.541 (0–6 m) | 10:15–10:35 | 14:30–17:30 | 14:30–17:30 |
|  |  |  | 0.261 (6–20 m) |  |  |  |
| Sep 18, 2018 | 6B | 124 | 1.029 | 9:35–10:00 | 14:15–17:15 | 14:15–17:15 |
| Oct 10, 2018 | 9B | 176 | 1.001 | 10:40–10:50 | 14:45–17:45 |  |
| Oct 25, 2018 | 9B | 1090 | 0.881 | 10:00–10:15 | 14:30–17:30 |  |
| Nov 16, 2018 | 12B | 901 | 1.198 (0–2 m) | 9:45–10:00 | 14:00–17:00 |  |
|  |  |  | 0.375 (2–20 m) |  |  |  |
| Dec 10, 2018 | 6B | 221 | 1.117 | 10:30–10:45 | 14:45–17:45 |  |
| Dec 19, 2018 | 17B | 255 | 0.451 (0–6 m) | 11:00–11:30 | 16:00–19:00 |  |
|  |  |  | 0.224 (6–20 m) |  |  |  |
| Jan 28, 2019 | 9B | 235 | 1.029 (0–2 m) | 10:30–10:45 | 14:30–17:30 |  |
|  |  |  | 0.387 (2–4.5 m) |  |  |  |
| Feb 7, 2019 | 12B | 279 | 0.536 (0–2 m) | 9:35–10:00 | 14:30–17:30 |  |

|  |  |  |  |  |  |
| --- | --- | --- | --- | --- | --- |
|  |  |  | 0.252 (2–20 m) |  |  |
| Apr 19, 019 | 9B | 445 | 1.916 (0–1 m) | 10:40–11:00 | 14:00–17:00 |
|  |  |  | 0.607 (1–5.5 m) |  |  |
| May 10, 2019 | 12B | 1711 | 0.894 (0–2 m) | 10:10–10:30 | 15:00–18:00 |
|  |  |  | 0.263 (2–20 m) |  |  |

### Sampling procedure

Vertical profiles of water temperature, pH, dissolved oxygen concentration, and specific conductivity were measured using a water quality sonde (EXO2; Xylem, Inc., Yellow Springs, OH, USA). At stations 12B and 17B, water samples were collected with a bucket at 0 m, with 10-L Niskin samplers at 2.5 m, and with 5-L Niskin bottles on a rosette sampler (AWS, JFE Advantech Co. Ltd., Kobe, Japan) at 5, 10, 15, and 20 m after *in situ* FRRf measurement (described below). Likewise, at stations 6B and 9B, water samples were collected with a bucket at 0 m, and with an electric pump at 2 and 3 m (6B) or 4 m (9B).

Macro-nutrients and dissolved inorganic carbon (DIC) concentrations were determined from an aliquot of 100 mL subsample collected at each depth, and immediately filtered through a syringe-type membrane filter (0.2 µm pore size, Acrodisc syringe filter; Pall Corporation, Ann Arbor, MI, USA) using clean techniques. The filtered samples were stored at −20°C until nitrate, nitrite, and phosphate analyses using an ion chromatograph system (Dionex Integrion HPIC system; Thermo Scientific, Waltham, MA, USA). DIC was analyzed using a total carbon analyzer (TOC-L; Shimadzu, Kyoto, Japan). For chlorophyll *a* (Chl-*a*) analysis, 50–200 mL sample was filtered onto a 25-mm glass-fiber filter (0.7 μm nominal pore size, GF/F; GE Healthcare, U.K. Inc., Little Chalfont, England). Chl-*a* was extracted with *N,N*-dimethylformamide for 24 h in the dark [61] and then stored at −80°C. The Chl-*a* concentration was determined with a 10-AU fluorometer (Turner Designs, Sunnyvale, CA, USA).

We measured underwater photosynthetically active radiation (PAR, 400–700 nm) from 30 m to the surface using a 2 π PAR sensor (CTG Ltd, West 106 Molesey, UK) along with the FRR fluorometer. We determined the diffuse attenuation coefficient *K*_d_ (m^−1^) with an exponential function as follows:

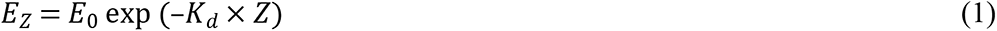

where *E*_Z_ and *E*_0_ are incident PAR (μmol photon m^−2^ s^−1^) at depth, *Z*, and 0 m, respectively. When the logarithmic slope of *K*_d_ significantly changed with depth, we calculated it for each layer (Table 2).

### FRRf measurements and photophysiological parameters

*In situ* induced Chl-*a* fluorescence profiles were measured vertically with a multi-excitation wavelength Fast Repetition Rate fluorometer (FRRf) system (FastOcean, S/N 17-0053-002; CTG Ltd, West 106 Molesey, UK). The field-type FRRf was equipped with two chambers for ambient light and dark readings. In order to remove ambient light noise, an optical bandpass filter (<670 nm) was attached above the light chamber. The dark chamber has black housing and piping with a pump to ensure that samples are measured under complete dark after 1–2 s of dark adaptation. Each chamber has three light-emitting diodes (LED) providing flash excitation energy centered at 444, 512, and 633 nm (S1 Appendix A). The 444 nm (blue) corresponds to the absorption peak of Chl-*a* while 512 nm (green) and 633 nm (orange) correspond to the absorption peaks of phycoerythrins and phycocyanins [49]. We employed four LED combinations to evaluate the green and orange excitation flashes: (1) 444 nm (2) 444 and 512 nm, (3) 444 and 633 nm, and (4) 444, 512 and 633 nm. We applied a single turnover method, which was consistent with a saturation phase (100 flashlets with 2 μs pitch) and a relaxation phase (40 flashlets with 50 μs pitch). This sequence was repeated eight times with a 100-ms interval for each LED combination. All combinations were repeated at least five times every 5 m from 20 to 10 m and every 2 m from 10 m to the surface at 12B and 17B, and every 0.5 m from the bottom to the surface layer at 6B and 9B during the up-cast, respectively. The power of flashlets (*E_FRRf_*) and the gain of the extra high tension of the photomultiplier tube (PMT eht) were optimized by FastPro8 software (version 1.0.50; CTG Ltd). All FRRf measurements were done between 09:30 and 11:30 (Table 2).

The concentration of PSII reaction center (RCII, nmol m^−3^) was estimated fluorometrically according to Oxborough et al. [28] as follows:

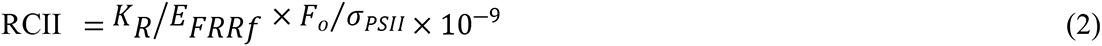

where *K_R_* is an instrument-specific constant (photons m^−3^ s^−1^), *F_o_* is the fluorescence intensity at the zeroth flashlet of a single turnover measurement when all RCII are open, and *σ_PSII_* is the absorption cross section of PSII photochemistry under dark (m^2^). A recent study showed *K_R_*/*E_FRRf_* can vary among phytoplankton taxonomic group and growth condition, and affect estimation of productivity [62]. In this study, we did not examine sample-specific *K_R_*/*E_FRRf_* values. Instead, we used a constant value as in Wei et al. [63], and taxonomic group and nutrients were assessed as the factors affecting Ф_e,C_ by statistical model approach (mentioned below).

The RCII-specific rate of electron transport based on FRRf (*J_f_,* μmol electrons nmol RCII^−1^ s^−1^) was calculated based on the Sigma Algorithm installed in FastPro8 [28,36]:

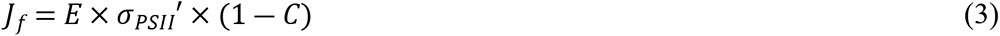

Where *E* is the incident PAR at each sampling depth (μmol photon m^−2^ s^−1^), *σ_PSII_*′ is the absorption cross section of PSII photochemistry under ambient light (m^2^), and (1 − *C*) is the fraction of RCII in the open state which is assumed, *qP* (=(*F′−F_O_′)/(F_m_′−F_O_′*)). Thus, the electron transport rate per water volume (*JV_f_,* μmol electrons m^−3^ s^−1^) was derived by *J_f_* × RCII

Phytoplankton can dissipate excessive energy as heat in PSII, and after PSII through alternative electron flows (AEFs) such as the Mehler reaction (for reviews, see [33,64]). Among these, the proportion of heat dissipation in total absorbed energy was estimated as the normalized Stern–Volmer quenching coefficient (NPQ_NSV_), which is equal to *F_O_′/F_v_′* [65]. This is commonly used to compare non-photochemical quenching among phytoplankton communities that have different light history [27,38,66]. Finally, the maximum quantum efficiency of PSII was evaluated by *F_v_/F_m_* [67].

### Evaluation of excitation wavelength combination

In order to assess the performance of the four wavelength combinations in the natural phytoplankton communities, we compared the minimum fluorescence yield *Fo* measured at each wavelength combination during cyanobacterial blooms (at 9B on August 28 in 2018), and during diatoms and zygnematophytes dominated (at 12 B on September 13 and at 6B on September 18 in 2018). We also compared electron transport rates estimated by FRRf (*JV_f_*, μmol e^−^ m^−3^ s^−1^) and by light-dark bottle method (*JV_O_*, μmol e^−^ m^−3^ s^−1^) [68] to verify the accuracy of estimated *JV_f_*.

The light-dark bottle method has been known to underestimate GPP due to differences in respiration rate between dark and light conditions [69,70], and oxygen oversaturation [71]. Nevertheless, since one molecule of CO_2_ fixation theoretically requires at least four electrons through photosynthesis, the method is still practical to determine whether the *JV_f_* determined by the FRRf was underestimated, or not. Water samples from each layer were poured into two or four 100-mL glass bottles. To measure the community respiration rate, another aliquot of sample water from each layer was poured into two 100-mL dark bottles. All bottles were incubated for 3 h in a growth-chamber (HCLP-880PF, Nippon Medical and Chemical Instruments Co., Ltd., Japan). Oxygen concentration of each bottle was measured by the optical oxygen spots and probe (Fibox 4; PreSens, Regensburg, Germany) before and after incubation. *JV_O_* was derived as

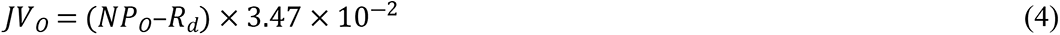

where *NP_O_* is net oxygen evolution rate (mg O_2_ m^−3^ h^−1^), *R_d_* is dark respiration rate (mg O_2_ m^−3^ h^−1^), and 3.47×10^−2^ is a conversion factor from hour to seconds, from mg O_2_ to μmol O_2_, and 4 mol e^−^ for 1 mol O_2_ evolution. Plots of *JV_O_* against light intensity were fitted in a two-parameter model as described in Webb et al. [72].

Additionally, quality control of all FRRf data measured with each excitation combination was assessed by the probability of an RCII being closed during the first flashlet of a single turnover saturation phase under dark (*Rσ_PSII_*) and ambient light (*Rσ_PSII_*′) by FastPro8 software. Although FastPro8 adjusted *Rσ_PSII_* and *Rσ_PSII_*′ around 0.05, these values changed dependent on depth, light environment and phytoplankton community composition. Thus, we compared the number of successful observations, *Rσ_PSII_* and *Rσ_PSII_* ′ among 4 combinations after rejection of extremely low-quality data (*Rσ_PSII_* or *Rσ_PSII_* ′ <0.03 or >0.08).

### Phytoplankton identification and enumeration

For enumerating phytoplankton, 50 mL of each sample was fixed with Lugol’s solution (1% of final concentration). After 24 h of settling in the dark, the supernatant was removed gently, and the sample was concentrated to 15 mL. Cells were counted under a light microscope at ×200 magnification where the size and volume of cells for each group were measured using cellSens software (Olympus, Tokyo, Japan) based on Hillebrand et al. [73]. All phytoplankton species were categorized into eight groups: diatoms, cyanophytes, small chlorophytes, zygnematophytes, cryptophytes, crysophytes, dinoflagellates, and euglenophytes. The phytoplankton community composition was assessed based on the carbon biomass converted from the biovolume [74].

### ^13^C uptake rate

Gross primary production per water mass (*GP_C_,* mg C m^−3^ h^−1^) was determined by a ^13^C-based method based on Hama et al. [16]. Water samples from each layer were taken to the laboratory and poured into two or four 500-mL polycarbonate bottles (Nalgene, Rochester, NY, USA), and spiked with NaH_13_CO_3_ to a final concentration of ca. 10% of ambient total inorganic carbon [75]. Incubations were initiated within 3 h after samples collected. Production experiments were conducted in incubators where the temperature and light environment were controlled in a growth-chamber along with an oxygen evolution experiment. Incubation PAR levels were set as follows: For site 12B and 17B, 100% and 65% of *E*_0_ for 0 m samples, and 30%, 10%, and 1.6% of *E*_0_ for 2.5-, 5- and 10-m samples; For site 6B, 100% and 65% of *E*_0_ for phytoplankton for the 0-m sample, 30% and 10% of *E*_0_ for the 2-m sample, and 1.6% of *E*_0_ for the 3-m sample; For site 9B, it was same as site 6B but 1.6% of *E*_0_ for the 4-m sample. Incubation PAR intensity was manipulated using black mesh filters covering polycarbonate bottles. Incubation temperature was set to the mean of respective sampling depths. The samples from 12B on 23 July and at 17B on 30 July were incubated under ambient light percentages (described above) and temperature on the balcony of Lake Biwa Environmental Research Institute (Shiga, Japan). Although incubation temperature was not controlled on 23 and 30 July, it changed <1.5°C during the incubations.

All incubations were conducted for 3 h. After each incubation period, water samples were filtered through pre-combusted (at 450°C for 4 h) 25-mm glass-fiber filters (0.7 μm nominal pore size, GF/F). The filters were stored at −20°C until final analysis. The carbon stable isotope ratio δ^13^C was measured using Delta V Advantage isotope ratio mass spectrometer coupled with Conflo IV interface and Flash 2000 elemental analyzer (Thermo Fisher Scientific, Waltham, MA USA) at the Isotope Research Institute (Yokohama, Japan). *GP_C_* was calculated according to Hama et all [16]. To compare with the measured *J_f_* in equation (3), *GP_C_* was converted to RCII-specific primary production rate (*PB_C_*, mg C nmol RCII^−1^ h^−1^) as follows:

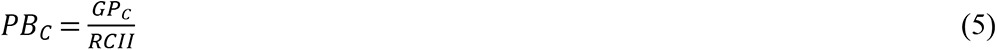

### Spectral correction

In order to account for the difference in spectral distribution and primary production response between ambient light field and artificial incubator light sources, we applied a spectral correction to the data-set to reduce any possible discrepancies between the methods. First, to correct the differences in the spectral distribution of excitation flash of FRRf and ambient light in water column, *σ_PSII_* was adjusted by spectral correction factor (SCF) following previous studies [27,34]:

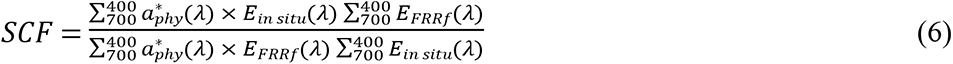

where 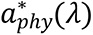 is the Chl-*a* specific absorption spectrum of phytoplankton (m^2^ mg Chl-*a*^−1^), *E_in situ_* (λ) and *E_FRRf_* (λ) are the spectral distribution of irradiance in the water column and excitation flash of FRRf, respectively. The 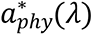, and *E_in situ_* (λ) were estimated from models described in previous studies where the spectral irradiance in the water column was estimated as follows [76,77],

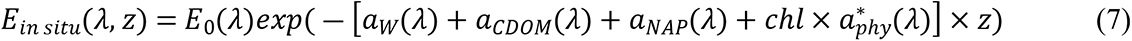

where *λ* is wavelength between 400 to 700 nm, *a_W_*, *a_CDOM_*, and *a_NAP_* are absorption spectrum of pure water, CDOM, and non-algal particles, respectively (m^−1^), and *chl* is Chl-*a* concentration (mg m^−3^) and z is depth (m). 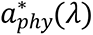 was estimated for each date and depth by Paavel’s model [78] for August, and Ylöstalo’s model [79] for other months in accordance with the species composition of phytoplankton community (S2 Appendix).

The *a_W_* was estimated by a common spectra model determined by Pope and Fry [80] (S3 Appendix A). The *a_CDOM_* was estimated by the equation *a_CDOM_* = *a_CDOM_* (320) exp(-*S*(λ-320)) [81]. We used previously measured, average values of limnetic sites from each basin, 1.03 and 2.28, for *a_CDOM (_*320) for the North Basin and the South Basin, and 0.017 for *S* [77,81] (S4 Appendix B). *a_NAP_* was estimated as *a_NAP_* = *a_NAP_* (440) EXP(-S (λ-440)) [82]. We used 0.264 for *a_NAP_* (440) and 0.004 for *S* as typical values for the area (S4 Appendix C). The calculated *E*_*in situ*_(*λ*, *z*) was adjusted with observed PAR intensity at each depth during each sampling date.

For spectral distribution of incident sunlight *E_0_* (*λ*) on each sampling date, we referred to the solar radiation spectrum database of central Japan in 2015 [83]. In order to fit the incident angle of sunlight and the weather condition on each sampling date, we used spectral data at 10 AM on January 30, February 9, April 8, May 17, July 2, 31, August 26, September 7, 19, October 10, 26, November 10, December 9 and 19 corresponding to each sampling date (S4 Appendix). For spectral irradiance during ^13^C incubation on July 23 and 30, we used spectral data at 4 PM on July 2 and 31 from the database.

Calculated SCFs are listed in the S1 Table in supporting information. The SCF for PAR intensity of each growth-chamber was also adjusted with the spectral distribution of the light source (S1 Appendix B) and 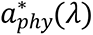 in the same manner as equation (6).

### Derivation of photosynthetic parameters

In order to calculate Ф_e,C_ (mol e^−^ mol C^−1^) from *J_f_* and *PB_C_* assessed at different light levels, photosynthesis versus irradiance curves (*P-E* curves) were obtained by curve-fitting using two *P-E* models. When *J_f_* or *PB_C_* showed photoinhibition (*i.e.*, decreasing *J_f_* or *PB_C_* with increasing *E* after the light-saturated phase), *P-E* curves were fitted in a three-parameter model as described in earlier studies [84–86]. When there was no photoinhibition, *P-E* curves were fitted in a two-parameter model as described in Webb et al. [72].

It should be noted that the phytoplankton community structures were occasionally different among the layers of water column, but the relationships between *J_f_* or *PB_C_* and *E* were well-fitted in the photosynthesis models, as in previous studies [25,30]. All parameters in the fitted models were calculated by function nls() in R ver. 3.4.3 [87]. The *PB_C_* values corresponding to underwater *E* of *in situ* FRRf measurements were extracted from the *P-E* curves. *PB_C_* can be functioned by *J_f_* with electron requirement for carbon fixation (Ф_e,C_) as

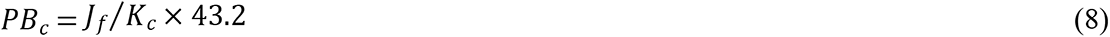

where 43.2 is the conversion factor from seconds to hour, and from μmol C to mg C, respectively.

### Data transformation and GLM for Ф_e,C_

Because FRRf measurements were inhibited by high light intensity (typically >1,000 μmol photons m^−2^ sec^−1^), presumably there were sampling biases between shallow and deep layers. To avoid this bias, we subsampled 240 observations from the data set on each sampling date by bootstrap sampling with replacements; 60 observations from four layers above the euphotic zone (0–3.75, 3.75– 7.5, 7.5–12.5, and 12.5–17.5 m) for the North Basin and 80 observations from three layers (0–1, 1–3 and 3–5.5 m) for the South Basin. Here a total of 3360 observations (1680 observations for the North Basin and 1680 observations for the South Basin) were used in all analyses of this study.

As in equation (8), Ф_e,C_ is defined as *J_f_* /*PBc* × 43.2. We modeled Ф_e,C_ using a generalized linear model (GLM) with gamma error distribution and a log-link function using the glm() function in R. We treated Ф_e,C_ as the dependent variable, and environmental factors as the explanatory variables. To avoid collinearity between the explanatory variables, Spearman’s *ρ* between all candidate factors were tested with a significance level of *p* < 0.05, and parameters with *ρ* ≥ 0.7 were regarded as collinear variables. We selected water temperature, PAR, turbidity, dissolved oxygen (DO), NH_4_, NO_3_ + NO_2_, PO_4_, *F_v_/F_m_*, *σ_PSII_*, Chl-*a*, and the fractions of diatom, cyanobacteria, and cryptophytes in phytoplankton community as the explanatory variables in GLM. Explanatory variables were standardized (mean 0 and standard deviation 1) after log-transformation. NH_4_ concentration and the fraction of phytoplankton group biomass lower than the detection limit were treated as 0.1 μmol L^−1^ and 0.1%, respectively. Multicollinearity of variables were further tested with the variance inflation factor (VIF) [88] using vif() in the package ‘car’ [89] in R. A set of all possible submodels was generated with the dredge() function in package ‘MuMIn’ [90], and the submodels were ranked based on the Akaike information criterion (AIC) [91].

## Results

### Evaluation of excitation wavelength combination

Vertical profiles of temperature showed weak stratification at Station 9B on August 28 and at Station 12B on September 13 (Fig. 2A, D) but not at Station 6B on September 18 (Fig. 2G). Chl-*a* concentration reached 42 μg L^−1^ at 2 m at Station 9B due to cyanobacterial bloom (Fig. 2C). Vertical profiles of the minimum PSII fluorescence yield (*Fo*) showed variability between four combinations of excitation wavelengths during cyanobacterial blooms (Fig. 2B), but not for diatom and zygnematophytes dominant communities (Fig. 2E, H). For example, *Fo* profiles derived by excitation lights at 444 nm and 444 + 512 nm were lower than excitation combinations at 444 + 633 nm and 444 + 512 + 633 nm when cyanobacteria were dominant at 0 and 2 m depth at Station 9B (South Basin) in August (Fig. 2B, C). On the other hand, there were no clear differences in *Fo* profiles at Station 12B (North Basin) on September 13 and Station 6B (South Basin) on September 18 when diatoms and zygnematophytes dominated (Fig. 2E, F, H, I). Further, the relationship between *JV_f_* and PAR intensity shows the utility of 633 nm for revealing signatures of cyanobacteria photosynthesis (Fig. 3). For example, *JV_f_* was clearly lower than *JV_O_* using excitation at 444 nm and 444 + 512 nm, but not at 444 + 633 nm and the combination of all three wavelengths at Station 9B on August 28 during a cyanobacteria bloom (Fig. 3A). No clear differences in *JV_f_* were observed between the combinations of wavelengths at Station 12B on September 13 and Station 6B on September 18, 2018 (Fig. 3B, C).

**Fig. 2.**
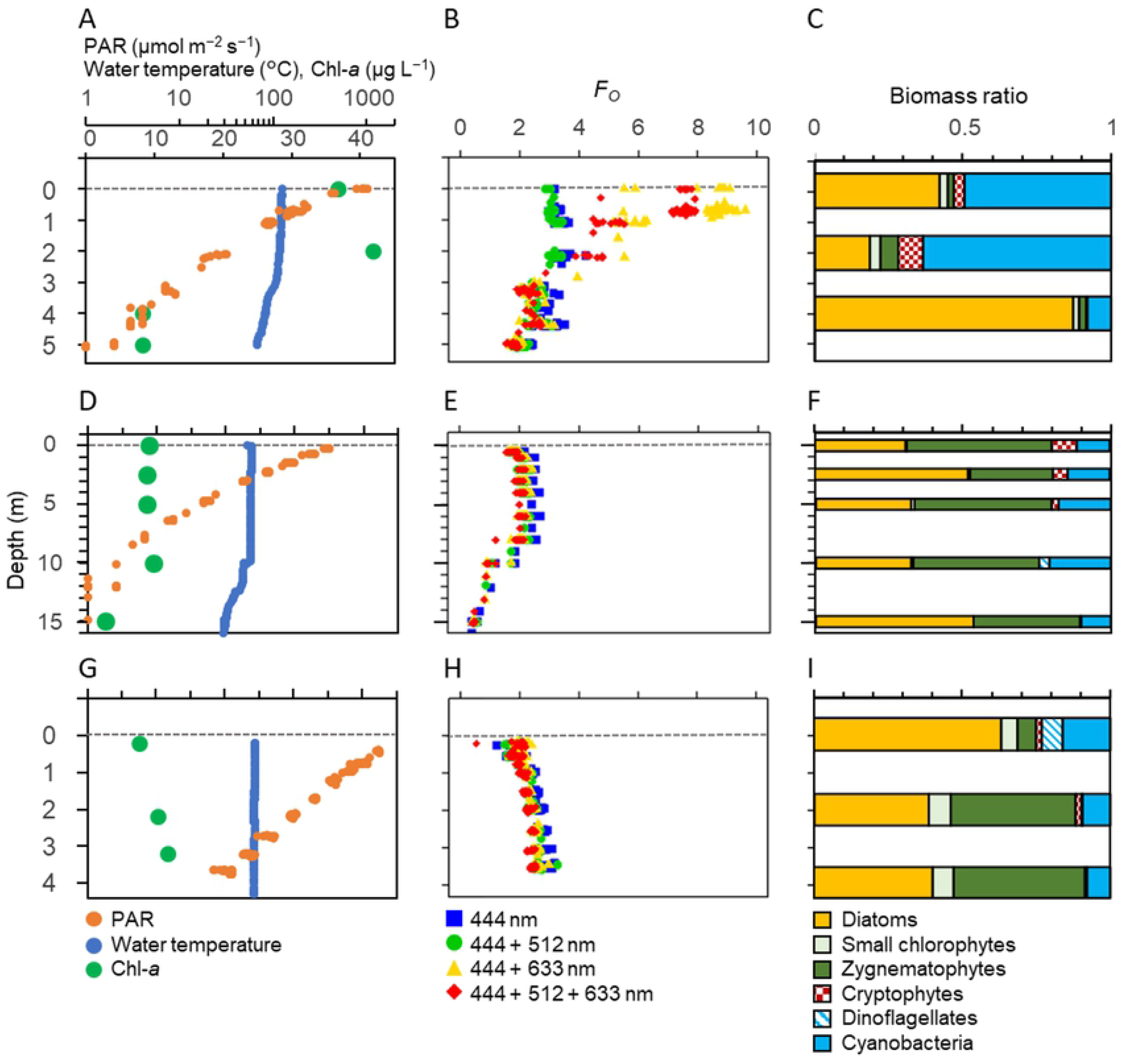
Vertical profiles of PAR, water temperature and Chl-*a* (A, D, G), *Fo* (B, E, H) estimated by different combinations of excitation wavelength from the FRRf, and phytoplankton biomass (C, F, I) at Station 9B in August 28 (A, B, C), Station 12B in September 13 (D, E, F) and Station 6B in September 18, 2018 (G, H, I). Grey dashed lines denote 0 m.

**Fig. 3.**
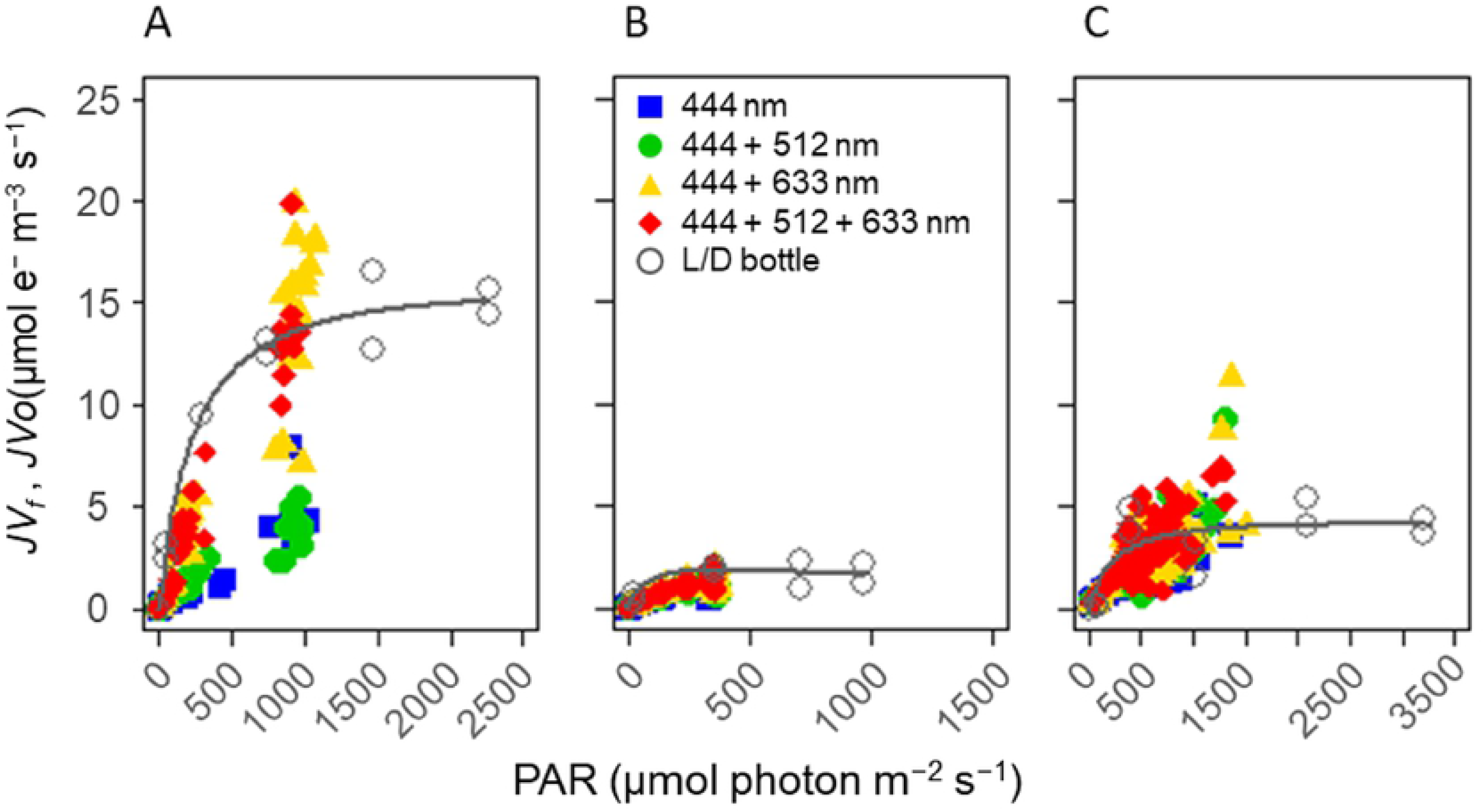
Scatter plots of *JV_f_* estimated by different combinations of excitation wavelength from the FRRf relative to ambient PAR intensity at (A) Station 9B on August 28, (B) Station 12B on September 13 and (C) Station 6B on September 18. The *JV_O_* estimates from the light-dark bottle method are also shown. For *JV_O_*, PAR intensity was calculated by light intensity of growth chambers and SCF (see Materials and methods). The fitted curve is given for *JV_O_* using a two-parameter model [72] to improve visibility.

The data quality of FRRf measurements during the study period showed reasonable stability (Table 3). Upon rejection of low-quality data (*e.g.*, with *Rσ_PSII_* or *Rσ_PSII_*′ < 0.03 or > 0.08), the number of successful observations was found to be highest when PSII was excited with a combination of three wavelengths, followed by excitation light at 444 + 633 nm, 444 + 512 nm, and 444 nm. Median values of both *Rσ_PSII_* and *Rσ_PSII_*′ were also near the optimal value (0.05) when the three wavelengths were combined.

**Table 3.**
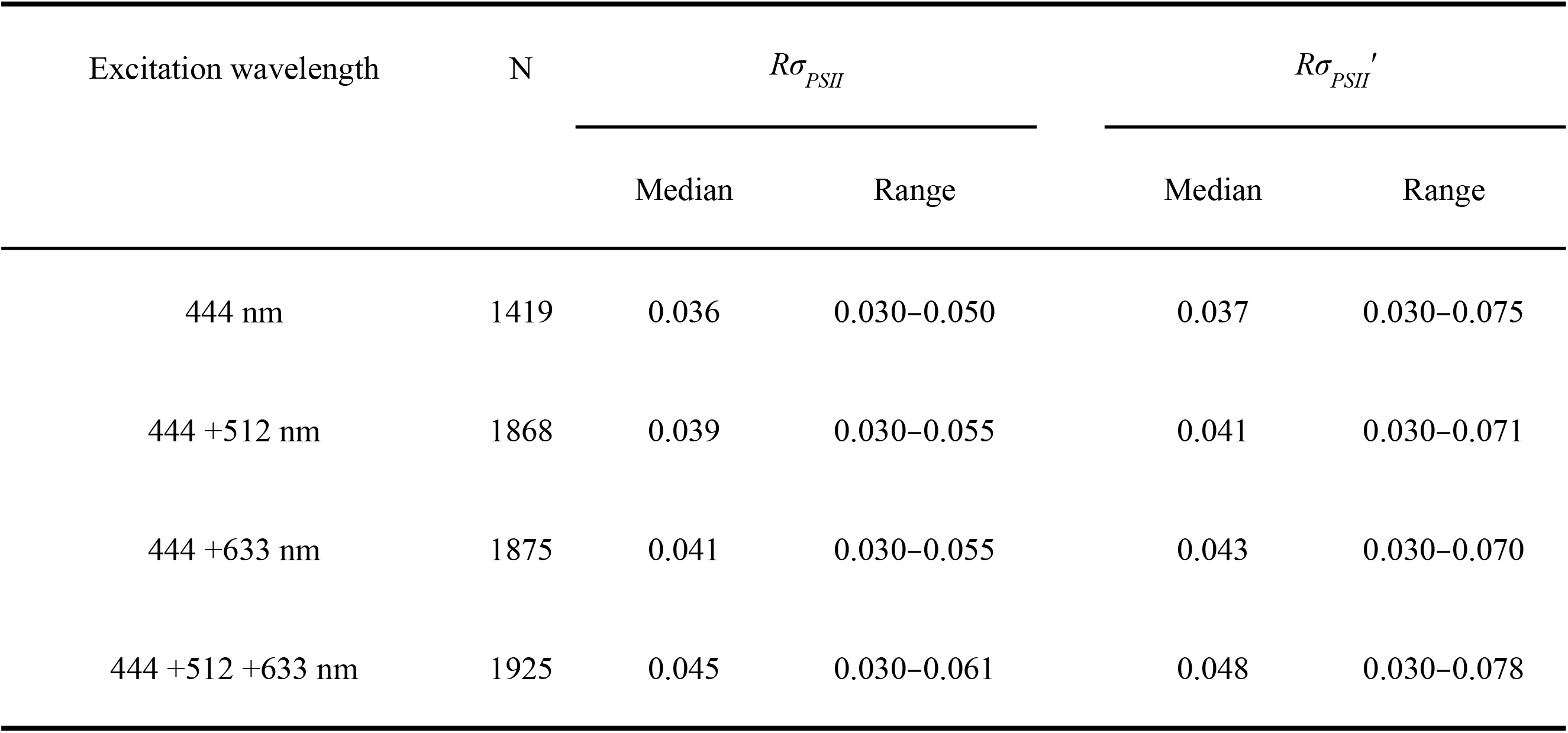
Comparison of the probability of RCII being closed during the first flashlet of a single turnover saturation phase under dark (*Rσ_PSII_*) and ambient light (*Rσ_PSII_*′) among four combinations of excitation wavelength. N, number of successful observations.

### Development of Ф_e,C_ model

#### Environmental and biological conditions

Ancillary measurements of water temperature, DO concentration, turbidity, Chl-*a*, NO_2_ + NO_3_, NH_4_ and PO_4_ concentrations from each sampling showed clear spatial and seasonal variability (Table 4). Water temperature varied from 7.5 to 30.2°C in the North Basin and 7.5 to 28.5°C in the South Basin throughout the study period (Table 4). NO_2_ + NO_3_ and NH_4_ concentrations were lower in summer and autumn, and higher in winter at both basins throughout the study period. PO_4_ did not show clear seasonal changes and was always lower than 0.04 μmol L^−1^ in both basins throughout the study period.

**Table 4.** Physical, chemical and biological (ancillary) conditions on sampling dates. The values denote depth ranges of 0 – 17.5 m for both Stations 17B and 12B, 0 – 4 m for Station 9B, and 0 – 3 m for Station 6B. NTU, nephelometric turbidity units; LOD, limit of detection.

| Date | Site | Temperature | DO | Turbidity | Chl-a | NO<sub>2</sub>+NO<sub>3</sub> | NH<sub>4</sub> | PO<sub>4</sub> |
| --- | --- | --- | --- | --- | --- | --- | --- | --- |
|  |  | (°C) | (%) | (NTU) | (µg L<sup>-1</sup>) | (µmol L<sup>-1</sup>) | (µmol L<sup>-1</sup>) | (µmol L<sup>-1</sup>) |
| Jul 23, 2018 | 12B | 17.8–30.2 | 82–116 | 0.3–1.3 | 2.4–13.0 | 0.1–1.1 | 0.2–0.4 | 0.01–0.03 |
| Jul 30, 2018 | 17B | 22.2–27.7 | 87–103 | 0.2–0.9 | 2.4–12.8 | 0.1–0.5 | 0.2–0.2 | 0.01–0.03 |
| Aug 28, 2018 | 9B | 25.0–28.5 | 79–136 | 1.6–8.2 | 8.0–41.9 | 0.4–1.1 | 0.8–1.1 | 0.02–0.02 |
| Sep 13, 2018 | 12B | 19.9–23.7 | 80–100 | 0.4–1.1 | 2.4–9.6 | 1.3–5.9 | 0.2–1.0 | 0.01–0.04 |
| Sep 18, 2018 | 6B | 24.2–24.4 | 102–103 | 2.2–4.3 | 7.6–12.4 | 0.3–1.7 | 1.0–1.2 | 0.01–0.02 |
| Oct 10, 2018 | 9B | 22.1–22.6 | 105–117 | 1.3–2.8 | 14.4–17.0 | 0.4–1.4 | 0.9–1.3 | 0.01–0.02 |
| Oct 25, 2018 | 9B | 19.6–19.8 | 107–110 | 1.0–1.9 | 16.2–17.3 | 0.1–0.4 | 0.8–0.8 | 0.01–0.03 |
| Nov 16, 2018 | 12B | 17.2–17.3 | 96–98 | 0.5–1.0 | 2.4–2.9 | 0.2–0.3 | 0.2–0.3 | 0.01–0.02 |
| Dec 10, 2018 | 6B | 11.5–11.6 | 98–99 | 2.2–2.7 | 7.6–8.0 | 0.1–0.5 | 0.8–0.9 | 0.03–0.03 |
| Dec 19, 2018 | 17B | 12.8–12.8 | 93–97 | 0.1–0.7 | 4.6–5.3 | 3.3–4.2 | 0.5–0.6 | 0.01–0.03 |
| Jan 28, 2019 | 9B | 7.5–8.3 | 96–98 | 1.1–2.6 | 2.5–3.4 | 4.8–5.9 | 1.0–1.2 | 0.02–0.03 |
| Feb 7, 2019 | 12B | 8.9–9.0 | 92–94 | 0.2–0.6 | 4.3–6.7 | 6.3–7.1 | 0.3–0.4 | 0.02–0.03 |
| Apr 19, 019 | 9B | 11.8–13.8 | 105–112 | 0.7–1.5 | 5.9–8.9 | 3.0–3.3 | <LOD | 0.01–0.03 |
| May 10, 2019 | 12B | 12.0–15.7 | 102–118 | 0.3–1.5 | 4.2–12.2 | 1.0–5.3 | <LOD–0.8 | 0.01–0.03 |

At all sampling dates, diatoms, zygnematophytes, cyanobacteria, and cryptophytes were the dominant groups in the phytoplankton biomass (S5 Appendix). Zygnematophytes, mainly composed of *Staurastrum dorsidentiferum*, *S. sebaldi*, and *Micrasterias hardyi,* were always found at all sites throughout the study period, except at Station 6B in December. Diatoms were dominant during summer to early autumn and reached 87% in the phytoplankton biomass at Station 6B in September (S5 Appendix). Cryptophytes were present at a relatively low proportion through the study period except at Station 6B in December. Cyanobacteria were mainly composed of *Anabaena* (*Dolichospermum*) *affinis* and *Aphanothece* sp. and bloomed at Station 9B in August 28. Small chlorophytes, crysophytes, and dinoflagellates always made up less than 20% of the total phytoplankton biomass. Euglenophytes were very rare and accounted for less than 0.5% of the total biomass through the study period.

#### Spatiotemporal variation and GLM development for Ф_e,C_

In order to develop an optimal electron requirement for carbon fixation (Ф_e,C_) model, we used the data set that was obtained by the combination of three excitation wavelengths due to the quality and reliability (Fig 2, 3). After bootstrap sampling, boxplots of median Ф_e,C_ were calculated for each sampling date (Fig. 4). Ф_e,C_ changed temporally from 1.1 to 31.0 mol e^−^ mol C^−1^ and were relatively higher in spring and summer in both the North and South basins. The mean annual Ф_e,C_ values were 5.6 mol e^−^ mol C^−1^ for the North Basin, 9.0 mol e^−^ mol C^−1^for the South Basin, and 7.3 mol e^−^ mol C^−1^ for all sampling sites.

**Fig. 4.**
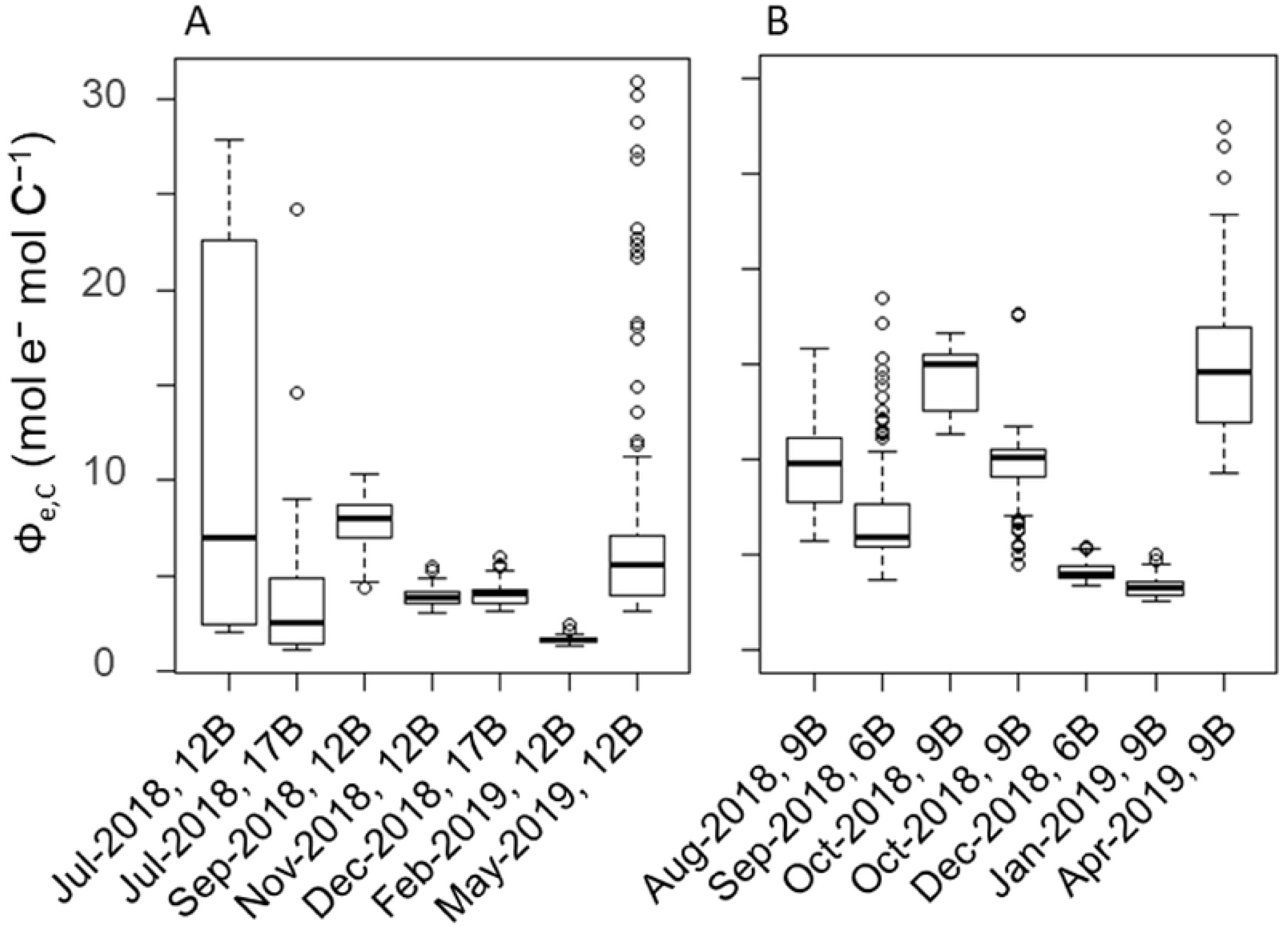
Spatial and temporal variability of Ф_e,C_ of the phytoplankton community in the North Basin (A) and the South Basin (B) through the study period. The box plot shows the median (bold line), the 25th (Q1) and the 75th (Q3) percentile. The whiskers indicate 1.5 times the interquartile range (Q3−Q1) below and above the Q1 and Q3. Outliers beyond the whiskers were plotted individually. Note: Ф_e,C_ values were derived from the data measured by the combination of three excitation wavelengths.

In order to select and define the explanatory variables for GLM, we examined correlations between Ф_e,C_ and all candidate environmental factors (Fig. 5). Small chlorophytes, crysophytes, and dinoflagellates were excluded because of their low proportion in the total phytoplankton biomass. Ф_e,C_ correlated positively with PAR, temperature, DO, NPQ_NSV_, Chl-*a*, and *σ_PSII_*, and negatively with maximum photochemical efficiency under dark conditions (*F_v_/F_m_*) throughout the study period. NPQ_NSV_ highly correlated with PAR and *F_v_/F_m_* (*ρ* = 0.70 and −0.96, respectively). RCII concentration highly correlated with Chl-*a* (*ρ* = 0.70), and diatoms and zygnematophytes also negatively correlated (*ρ* = −0.70) with each other. Based on the correlation matrix, we selected temperature, PAR, turbidity, DO, *F_v_/F_m_*, *σ_PSII_*, NH_4_, NO_2_ + NO_3_, PO_4_, Chl-*a*, and fractions of diatoms, cyanobacteria, and cryptophytes in the phytoplankton biomass as explanatory variables for the GLM. The effects of PAR and diatoms in this analysis may have included those of NPQ_NSV_ and zygnematphytes.

**Fig. 5.**
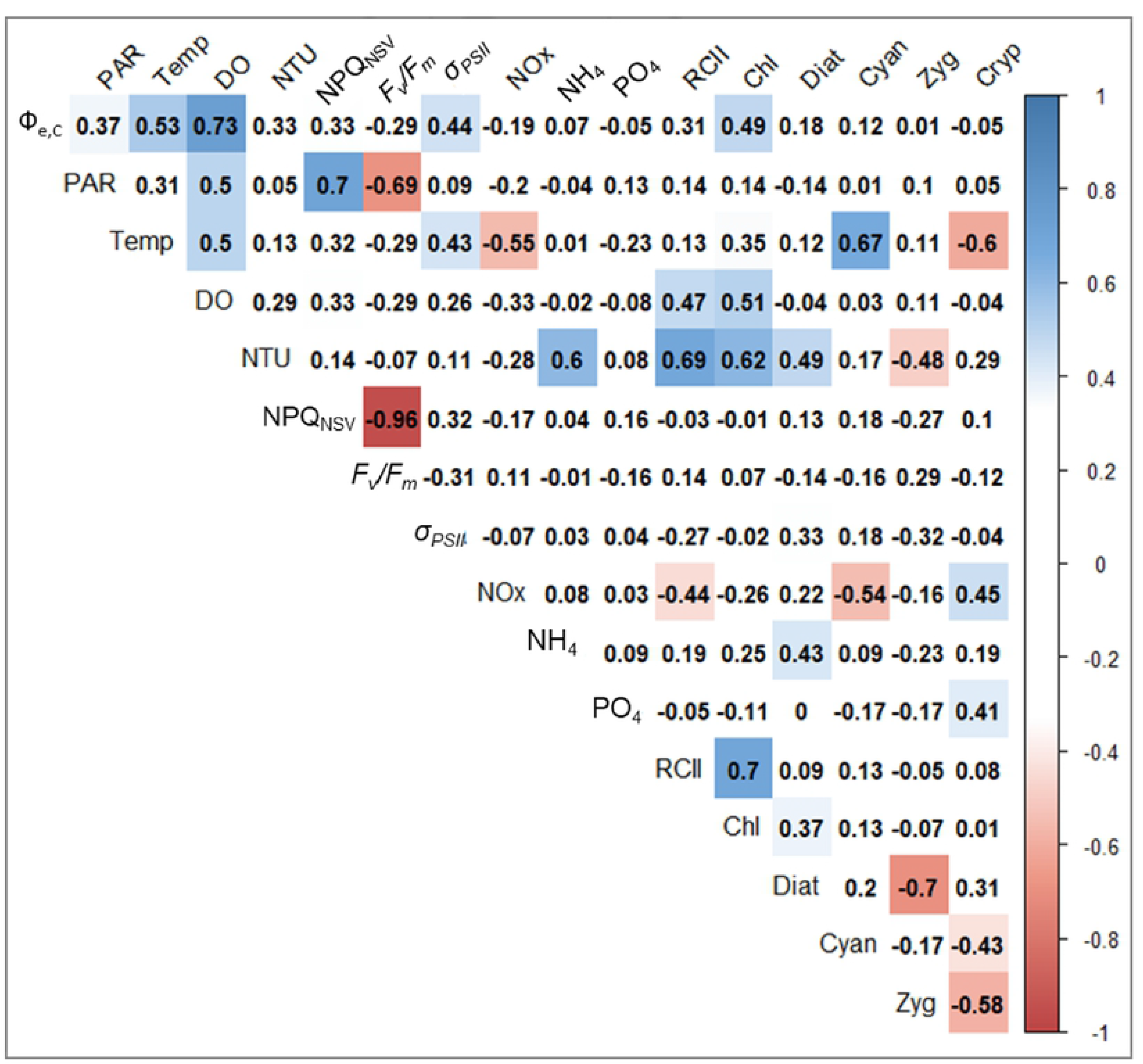
Matrix of Spearman’s *ρ* between photosynthetic parameters measured by excitation wavelengths of 444 + 512 + 633 nm, physicochemical factors, and biomass fraction of each phytoplankton group. Colored panels denote statistical significance (*p* < 0.05). Abbreviations of variables are: Temp, water temperature; NTU, turbidity; Chl, Chl-*a* concentration; Diat, diatom; Cyano, cyanobacteria; Zyg, zygnematophytes; Cryp, cryptophytes.

Among all possible models, the best model with the lowest AIC was the full model without PAR (Table 5). All variables in the best model exhibited VIF < 10, and thus, collinearity was negligible. The *R*^2^ for the best model was 0.67. Among the explanatory variables, temperature showed the highest significance in the best model (coefficient of 0.51), followed by cyanobacteria (coefficient of −0.20), and *σ_PSII_* (coefficient of 0.17). The performance of more parsimonious models were examined to evaluate the laborious sampling effort of nutrients and microscopy analysis phytoplankton assemblages. The lowest AIC models without nutrients (Model 2), and without nutrients and phytoplankton assemblages (Model 3) were employed. The Model 2 included six variables, (temperature, *F_v_/F_m_*, *σ_PSII_*, cyanobacteria, diatoms and cryptophytes), while Model 3 included three variables (temperature, *F_v_/F_m_*, and *σ_PSII_*). The values for *R*^2^ for the Model 2 and the Model 3 were 0.61 and 0.42, respectively. The results of the other sub-models with and without standardization of variables are listed in the supporting information (S2 Table and S3 Table).

**Table 5.**
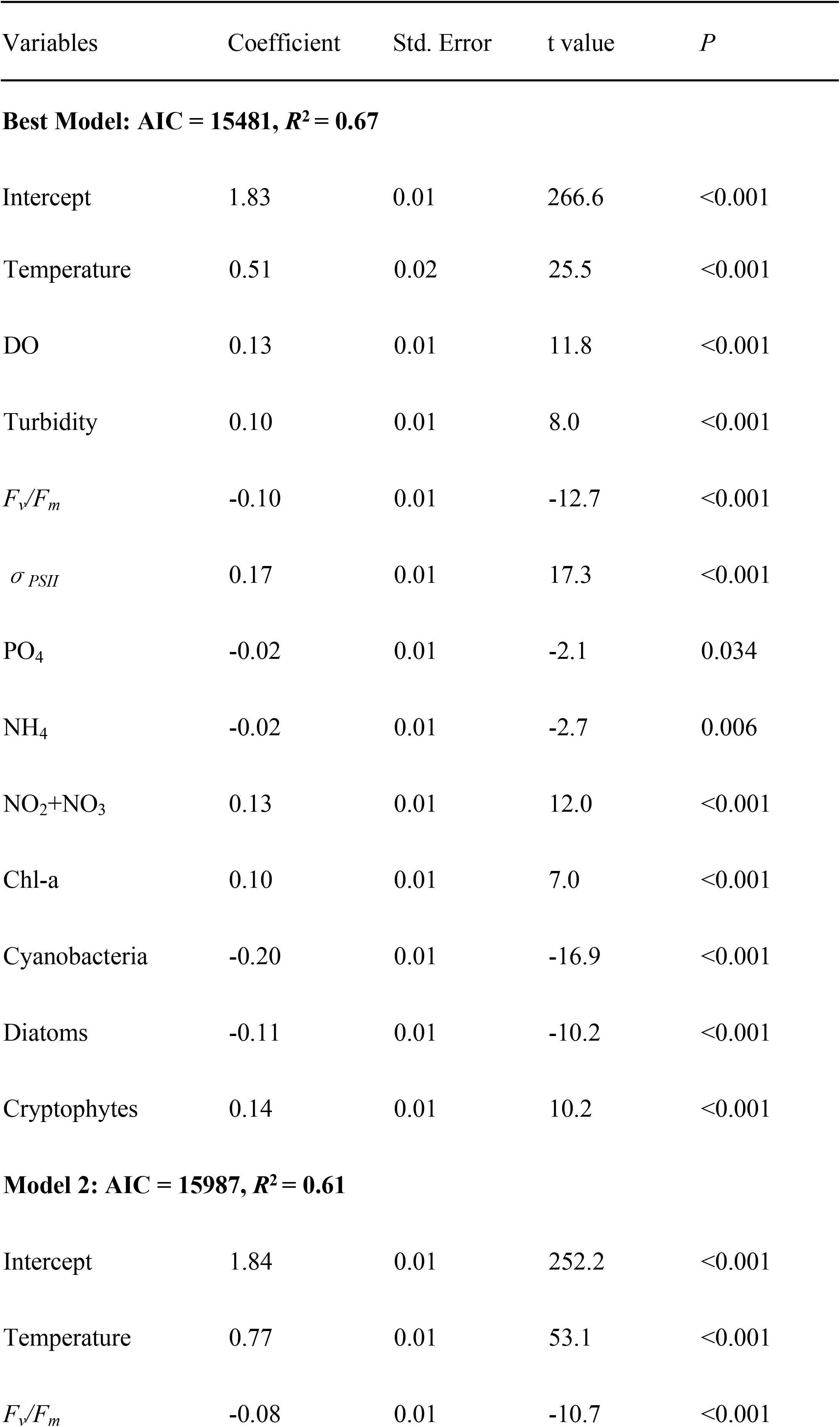
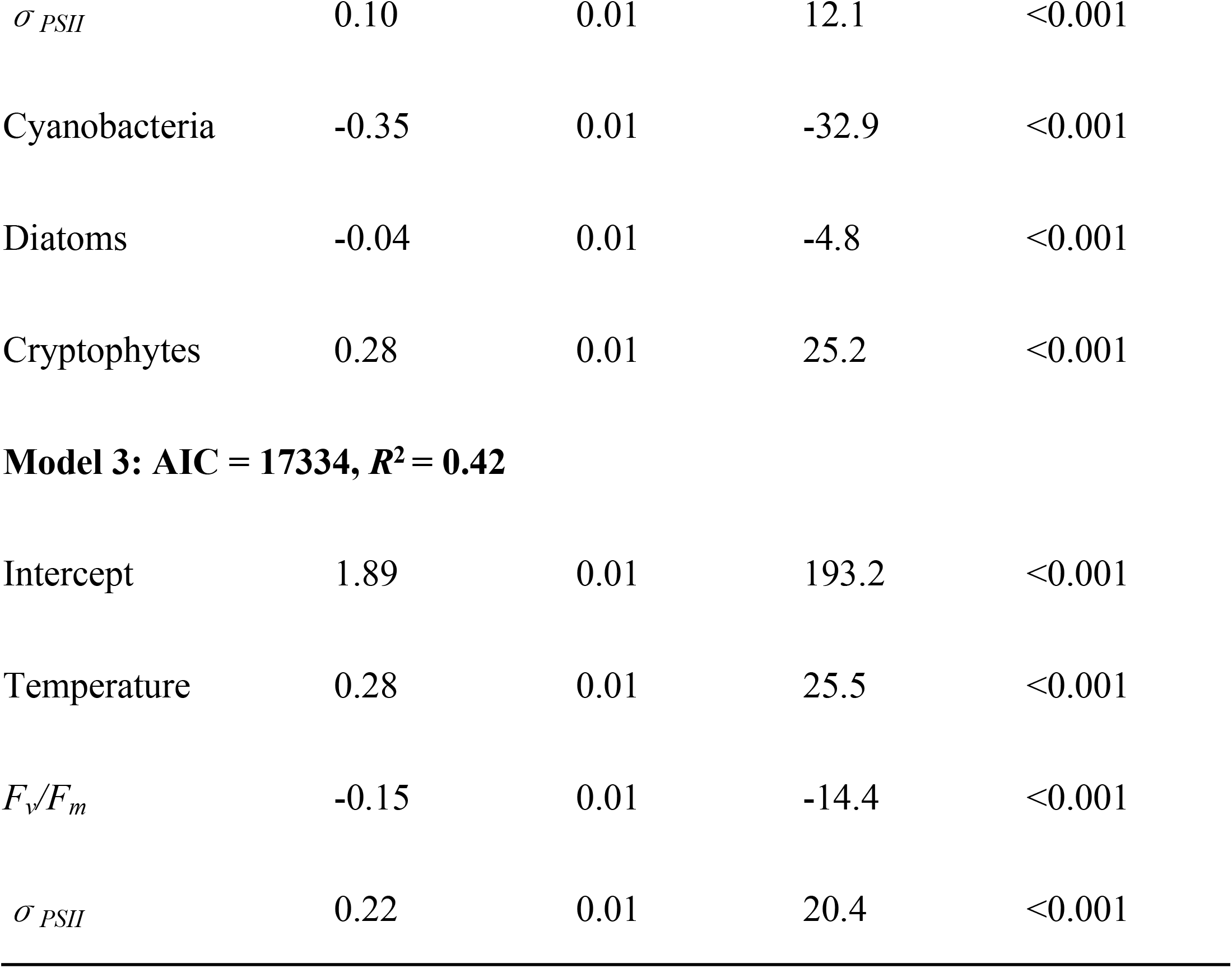
Statistical results of the GLM analysis showing the best model (smallest AIC in all models), the Model 2, and the Model 3 for Ф_e,C_. Coefficients were derived for log-transformed and standardized variables (see Materials and methods). AIC and *R*^2^ for each model are also shown.

| Variables | Coefficient | Std. Error | t value | *P* |
| --- | --- | --- | --- | --- |
| **Best Model: AIC = 15481, $R^2$ = 0.67** |  |  |  |  |
| Intercept | 1.83 | 0.01 | 266.6 | <0.001 |
| Temperature | 0.51 | 0.02 | 25.5 | <0.001 |
| DO | 0.13 | 0.01 | 11.8 | <0.001 |
| Turbidity | 0.10 | 0.01 | 8.0 | <0.001 |
| $F_v/F_m$ | -0.10 | 0.01 | -12.7 | <0.001 |
| $\sigma_{PSII}$ | 0.17 | 0.01 | 17.3 | <0.001 |
| PO<sub>4</sub> | -0.02 | 0.01 | -2.1 | 0.034 |
| NH<sub>4</sub> | -0.02 | 0.01 | -2.7 | 0.006 |
| NO<sub>2</sub>+NO<sub>3</sub> | 0.13 | 0.01 | 12.0 | <0.001 |
| Chl-a | 0.10 | 0.01 | 7.0 | <0.001 |
| Cyanobacteria | -0.20 | 0.01 | -16.9 | <0.001 |
| Diatoms | -0.11 | 0.01 | -10.2 | <0.001 |
| Cryptophytes | 0.14 | 0.01 | 10.2 | <0.001 |
| **Model 2: AIC = 15987, $R^2$ = 0.61** |  |  |  |  |
| Intercept | 1.84 | 0.01 | 252.2 | <0.001 |
| Temperature | 0.77 | 0.01 | 53.1 | <0.001 |
| $F_v/F_m$ | -0.08 | 0.01 | -10.7 | <0.001 |
| $\sigma_{PSII}$ | 0.10 | 0.01 | 12.1 | <0.001 |
| Cyanobacteria | -0.35 | 0.01 | -32.9 | <0.001 |
| Diatoms | -0.04 | 0.01 | -4.8 | <0.001 |
| Cryptophytes | 0.28 | 0.01 | 25.2 | <0.001 |
| **Model 3: AIC = 17334, $R^2 = 0.42$** |  |  |  |  |
| Intercept | 1.89 | 0.01 | 193.2 | <0.001 |
| Temperature | 0.28 | 0.01 | 25.5 | <0.001 |
| $F_v/F_m$ | -0.15 | 0.01 | -14.4 | <0.001 |
| $\sigma_{PSII}$ | 0.22 | 0.01 | 20.4 | <0.001 |

## Discussion

Recently, studies using FRRf with only one wavelength, around 450 nm, for primary production measurements have been successful [25,26]. However, the absorption spectrum of phytoplankton is highly dependent on the construction of antenna pigments, and is quite different between cyanobacteria and the other phytoplankton groups [92]. The present study demonstrated that *JV_f_* measured with an excitation light of 444 nm (single source), or with excitation lights of 444 + 512 nm was considerably underestimated compared to measurements utilizing 633 nm, particularly when cyanobacteria dominated (Fig. 3). This underestimation is expected because of the mismatch in wavelength between excitation wavelengths of the FRRf and the absorption spectrum of cyanobacteria. The blue excitation flash at 444 nm can fail to saturate the RCII in cyanobacteria during a single turnover measurement of FRRf [41,93], and thus underestimate *Fo* [94] and GPP [21]. Although we estimated *JV_f_* by each combination of excitation wavelength with SCF, which was calculated from extrapolated 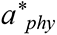, there might be large differences between modeled and actual 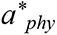 around 633 nm, but not around 512 nm in the green spectrum. Indeed, 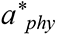 at 630 nm in August, when *Anabaena* spp. dominated, was estimated as 0.008−0.009 m^-2^ mg Chl-*a*^-1^ (S2 Appendix) while that of cultured *Anabaena* sp. was 0.028 m^-2^ mg Chl-*a*^-1^ [49]. Further, 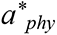 of cyanobacteria can vary with taxonomic group [49] and nutrient availability [95,96]. Our results suggest that if the absorption spectrum cannot be measured *in situ*, the excitation light targeting phycobilin antenna pigments should be used to measure ETR_PSII_ in cyanobacterial dominant communities.

In this study, Ф_e,C_ in Lake Biwa ranged temporally from 1.1 to 31.0 mol e^−^ mol C^−1^ (Fig. 4). Although our Ф_e,C_ range varied less than those reported in temperate ocean conditions (1.0 to 66.5 mol e^−^ mol C^−1^ [26]), it was similar to that in an Atlantic Ocean transect (1.1 to 28.2 mol e^−^ mol C^−1^ [35]) and Ariake Bay (1.2 to 26.6 mol e^−^ mol C^−1^ [25]), and those in a shallow-eutrophic lake (from 14.7 to 38.6 mol e^−^ mol C^−1^ [58]). Considering that environmental conditions varied substantially among the sampling sites and seasons in this study, the range of our Ф_e,C_ reasonably represented variations in Lake Biwa, including oligotrophic (North Basin) and mesotrophic (South Basin) areas.

The GLM determining Ф_e,C_ revealed that multiple physicochemical and biological factors, except PAR, were significant in Lake Biwa (Table 6). Previous studies primarily focused on the relationships between light environment, or NPQ_NSV_ and Ф_e,C_ [25,27,34,38,40,63]. The NPQ_NSV_ is mechanistically linked with alternative electron flow (AEF) activity, which is activated by excess light and photodamage on PSII [1,64,97]. In the present study, PAR intensity was highly correlated with NPQ_NSV_ (Fig. 5), and thus we incorporated PAR in GLM for Ф_e,C_ as a proxy of excess light and NPQ_NSV_. Although PAR was not utilized in the development of the best model, water temperature, DO, *F_v_/F_m_, σ_PSII_*, nutrient conditions, and the compositions of the phytoplankton community were selected. Our results suggest that the light environment is not always the primary factor determining Ф_e,C_, rather, water temperature plays a critical role in the electron requirement for carbon fixation. Generally, increased temperature decreases the CO_2_ affinity of Rubisco through the acceleration of the O_2_ evolution rate, and reduction of O_2_ solubility [98,99]. Furthermore, increasing temperature may lead to a state of chronic photoinhibition through photodamage [100], as well as changes in species composition of the community [40]. Although we did not examine the interaction among all explanatory variables, temperature may have affected Ф_e,C_ *vis-à-vis* interaction with the other factors such as nutrient stoichiometry [101].

The relationships between multiple environmental factors other than light and Ф_e,C_ were reported in previous FRRf [35,36,40] and PAM studies [102,103]. Lawrenz et al. [35] investigated the relationships between the Ф_e,C_ of marine phytoplankton and environmental conditions, and explored the best fitting models with 14 data sets from various geographic regions. They examined water temperature, salinity, optical depth, attenuation coefficient, Chl-*a*, NO_3_, and PO_4_ as explanatory variables, and showed only water temperature, NO_3_, and PO_4_ significantly correlated with Ф_e,C_ in the data set. Their results support our best model which included water temperature and nutrients. However, Lawrenz et al. also showed that the coefficients and significance of the identified parameters are quite dependent on region of interest. Thus, future studies are necessary to examine the complex relationship between temperature, light and nutrients conditions in freshwater environments to delineate specific conditions that explain Ф_e,C_ discrepancies.

Although three excitation wavelength combinations were used to evaluate cyanobacterial photosynthesis correctly, the proportion of cyanobacteria in phytoplankton biomass was significant in determining the best Ф_e,C_ model (Table 5). The proportion of cyanobacteria within the community may cause differences in the light absorption characteristics between cyanobacteria and the other phytoplankton groups; cyanobacteria absorb only 25–30% of light in PSII, while other phytoplankton absorb 48–58% [42]. Kromkamp et al. (2008) suggested that most of the Chl-*a* in cyanobacteria is associated with PSI rather than PSII, and detection-limited by fluorometry. The FRRf method primarily targets pigments associated with PSII, thus the electron flow initiated by the excited PSI is difficult to evaluate. Therefore, evaluating the proportion of cyanobacteria in phytoplankton biomass might be crucial for correcting GPP estimation with ETR_PSII_ in natural phytoplankton communities.

As expected, PO_4_ concentration negatively affected Ф_e,C_ (Table 5). This result was consistent with previous PO_4_ manipulation experiments conducted with marine phytoplankton [104]. The effect of PO_4_ was, however, one of the lowest among all factors (Table 5). The low significance is likely due to the fact that the PO_4_ concentrations did not change drastically and were generally low (0.01–0.04 μmol L^−1^, Table 4) throughout the study period.

In order to verify the performance of the Ф_e,C_ models, we subsequently compared daily GPP estimated with FRRf (*GPP_f_*) and ^13^C (*GPP_13C_*) for the North and the South basins on each sampling date. The RCII-specific primary production based on FRRf (*PB_f_*, mg C mmol RCII ^−1^ h^−1^) was calculated from the *J_f_*, and computed Ф_e,C_ with the best model, Model 2 and Model 3. The relationships between *PB_f_* and PAR were fitted in *P-E* curves with models [72,84–86], as described in the materials and methods. Finally, daily *GPP_f_* and *GPP_13C_* were estimated as follows:

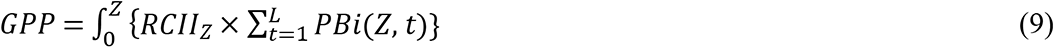

where *RCII*_*Z*_ is the RCII concentration at depth *Z*, *L* is the day length (h), and *PBi*(*Z*, *t*) is *PB_f_* or *PB_13C_* at depth *Z* and time *t* (h). The *RCII*_*Z*_ and *PBi*(*Z*, *t*) were calculated every 1.25 m for the North Basin and every 0.5 m for the South Basin based on average values of the observed data. Day length and daily PAR data were obtained from the Japan Meteorological Agency for the North Basin, and were measured by a PAR sensor (PAR-02D; Prede Co., ltd., Tokyo, Japan) at Otsu for the South Basin.

The estimated *GPP_13C_* varied from 71 to 787 g C m^−2^ d^−1^, while *GPP_f_* with best model, the Model 2 and the Model 3 varied from 86 to 630 g C m^−2^ d^−1^, 85 to 729 g C m^−2^ d^−1^ and 117 to 906 g C m^−2^ d^−1^, respectively (for date-specific values, see S4 Table). The *GPP_f_* using the best model relative to *GPP_13C_* varied from 0.48 to 1.46, suggesting that FRRf parameters with the best Ф_e,C_ model can reasonably reproduce *GPP_13C_* in Lake Biwa (Fig. 6). Relative *GPP_f_* with Model 2 and Model 3, including fewer variables, also replicated *GPP_13C_* well, varying from 0.46 to 1.59 and 0.47 to 2.11, respectively. The comparative results suggest that even if nutrients and phytoplankton biomass are not considered, *GPP_13C_* can be estimated using the FRRf parameters coupled with temperature for both oligotrophic and mesotrophic area in Lake Biwa. This model parameterization is significant for future applications in freshwater ecosystems where environmental conditions and phytoplankton communities can vary spatially and temporally.

**Fig. 6.**
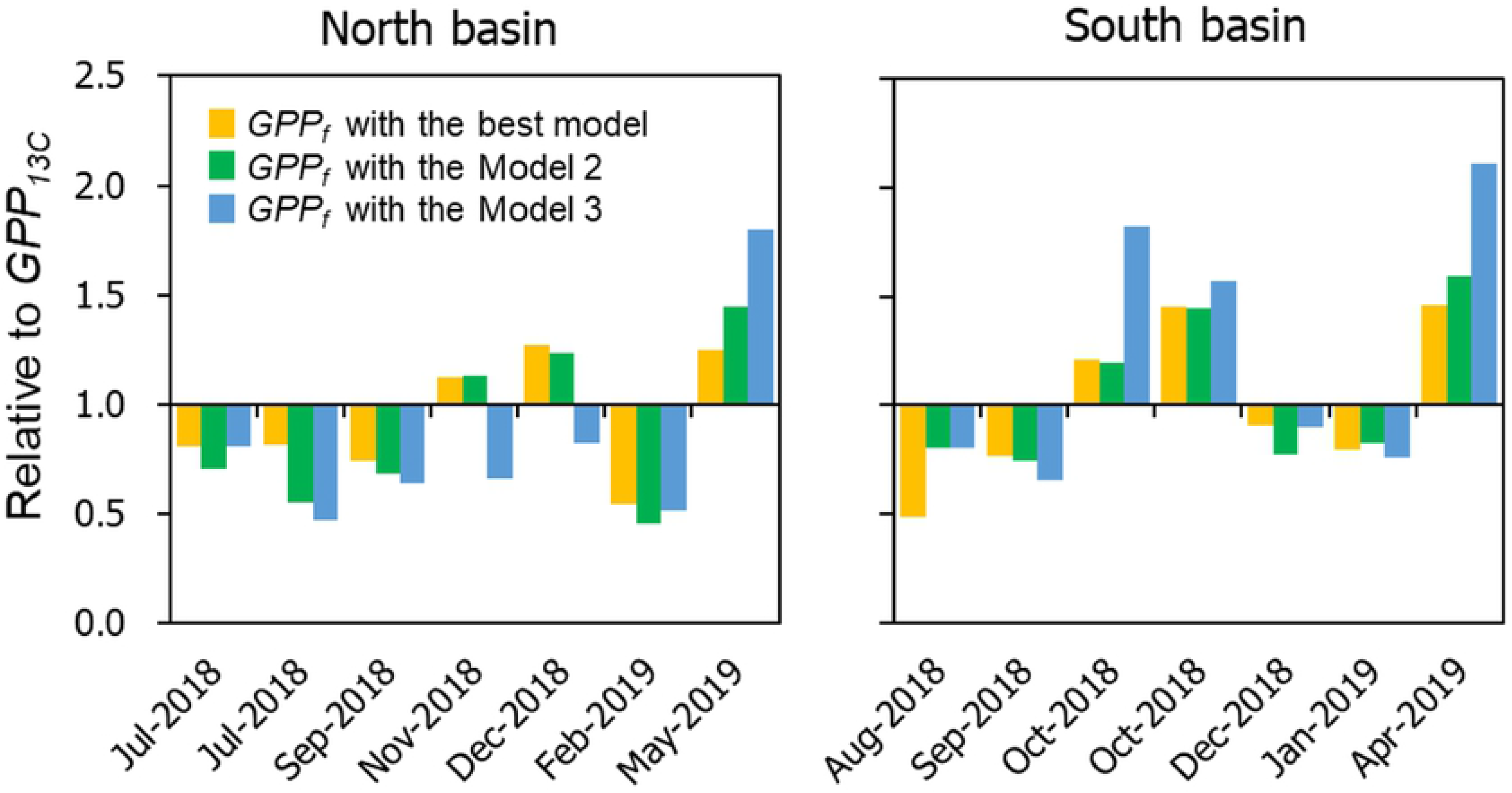
Relative *GPP_f_* estimated by ETR_PSII_ with the Ф_e,C_ models against the *GPP_13C_* in each sampling date for North and South basins.

In conclusion, the FRRf equipped with excitation wavelengths 633 nm is effective in estimating ETR_PSII_ for freshwater phytoplankton communities during cyanobacterial blooms. Contrary to our hypothesis, water temperature was the most important determinant factor, while phosphorus concentration was less effective in the Ф_e,C_ model in Lake Biwa. Further, *GPP_13C_* dynamics were effectively estimated from the ETR_PSII_ using the best Ф_e,C_ model for both oligotrophic and mesotrophic areas in Lake Biwa. The parsimonious model, including only temperature and two photosynthetic parameters also sufficiently reproduced *GPP_13C_*. In this study, the phytoplankton community was verified through microscopy, but fluorometric characteristics from the FRRf may also allow researchers to determine phytoplankton assemblages at high *in situ* spatial and temporal resolution [105], and thus simplify the GPP measurements during cyanobacterial blooms. This study provides strong validation for measurements of primary productivity by FRRf in lakes with large spatiotemporal variabilities of phytoplankton assemblages and environmental conditions from oligo- to mesotrophic lakes (or lacustrine) environments. In the future, bio-optical measurements will allow researchers to disentangle the causality between anthropogenic nutrient control and fish catch [13,59,106], and between climate change and the production of higher trophic levels [7,107].

## Acknowledgments

We would like to thank Dr. Takamaru Nagata in LBERI and Dr. Hiroki Haga in Lake Biwa Museum for sharing their ship time and valuable support during sampling. We sincerely thank Dr. Satoshi Nakada and Maho Iwaki for providing PAR profile data, and Hirokazu Teraishi for assisting with the chemical analysis. This study was supported by the Collaborative Research Fund from Shiga Prefecture entitled “Study on water quality and lake-bottom environment for protection of the soundness of water environment” under the Japanese Grant for Regional Revitalization, and the Environment Research and Technology Development Fund (No. 5-1607) of the Ministry of the Environment, Japan.

## Author contributions

Conceived and designed the experiments: TK KK. Performed the experiments: TK KH KS. Analyzed the data: TK. Contributed reagents/materials/analysis tools: KH. Supervision: AI. Wrote the original draft: TK. Review and Editing: KH KS VSK AI KK.

---

**S1 Table. Spectral correction factor (SFC) for *JV_f_* estimation in each sampling date.**

**S2 Table. All GLM submodels with standardization of variables.**

**S3 Table. All GLM submodels without standardization of variables**

**S4 Table. GPP (g C m^−2^ d^−1^) estimated by ^13^C and FRRf with Ф_e,C_ models in each sampling date.**

**S1 Appendix. Spectral distribution of (A) excitation flash of FastOcean (B) light source of growth-chamber.**

**S2 Appendix. Modeled absorption spectrum for various Chl-*a* concentration.** The spectrum were calculated with Paavel’s model [78] for 30 and 40 μg L^−1^ in August, and Ylöstalo’s model [79] for others.

**S3 Appendix. Modeled absorption spectrum of (A) pure water, (B) CDOM and (C) non-algal particles.**

**S4 Appendix. Spectral distribution of incident sunlight at 10:00 in (A) April to September, (B) October to February and (C) at 4 PM in July in 2015.** Each spectrum distribution was referred to calculate the spectral correction in each sampling date as in equation (7).

**S5 Appendix. Relative contribution to total phytoplankton biomass by algal groups on each sampling date.**

